# Antibodies generated *in vitro* and *in vivo* elucidate design of a thermostable ADDomer COVID-19 nasal nanoparticle vaccine

**DOI:** 10.1101/2023.03.17.533092

**Authors:** Dora Buzas, H Adrian Bunzel, Oskar Staufer, Emily J Milodowski, Grace L Edmunds, Joshua C Bufton, Beatriz V Vidana Mateo, Sathish K N Yadav, Kapil Gupta, Charlotte Fletcher, Maia Kavanagh Williamson, Alexandra Harrison, Ufuk Borucu, Julien Capin, Ore Francis, Georgia Balchin, Sophie Hall, Mirella Vivoli Vega, Fabien Durbesson, Renaud Vincentelli, Joe Roe, Linda Wooldridge, Rachel Burt, J L Ross Anderson, Adrian J Mulholland, Bristol UNCOVER Group, Jonathan Hare, Mick Bailey, Andrew D Davidson, Adam Finn, David Morgan, Jamie Mann, Joachim Spatz, Frederic Garzoni, Christiane Schaffitzel, Imre Berger

## Abstract

COVID-19 continues to damage populations, communities and economies worldwide. Vaccines have reduced COVID-19-related hospitalisations and deaths, primarily in developed countries. Persisting infection rates, and highly transmissible SARS-CoV-2 Variants of Concern (VOCs) causing repeat and breakthrough infections, underscore the ongoing need for new treatments to achieve a global solution. Based on ADDomer, a self-assembling protein nanoparticle scaffold, we created ADDoCoV, a thermostable COVID-19 candidate vaccine displaying multiple copies of a SARS-CoV-2 receptor binding motif (RBM)-derived epitope. *In vitro* generated neutralising nanobodies combined with molecular dynamics (MD) simulations and electron cryo-microscopy (cryo-EM) established authenticity and accessibility of the epitopes displayed. A Gigabody comprising multimerized nanobodies prevented SARS-CoV-2 virion attachment with picomolar EC_50_. Antibodies generated by immunising mice cross-reacted with VOCs including Delta and Omicron. Our study elucidates nasal administration of ADDomer-based nanoparticles for active and passive immunisation against SARS-CoV-2 and provides a blueprint for designing nanoparticle reagents to combat respiratory viral infections.

## Introduction

As of January 2023, the COVID-19 pandemic, caused by severe acquired respiratory syndrome coronavirus 2 (SARS-CoV-2), continues to spread globally, with over 660 million confirmed cases and close to 7 million deaths reported worldwide according to the World Health Organisation (WHO). The economic damage caused by the pandemic has been significant, with many countries having experienced severe economic downturns as a result of lockdowns and other non-pharmaceutical intervention measures implemented to slow the spread of the virus. In an unprecedented effort, a large number of COVID-19 vaccine candidates were developed at record speed ^1–4^, and several were authorised for emergency use or received full approval by regulatory agencies around the world ^1^. Among these, prominent examples include the Pfizer-BioNTech (Cominatry) and Moderna (Spikevax) vaccines which use lipid-encapsulated messenger RNA (mRNA) to instruct cells to produce the SARS-CoV-2 spike glycoprotein (S) to trigger an immune response ^5–10^. The Oxford-AstraZeneca ((Vaxzevria) and Johnson & Johnson (Jcovden) vaccines use adenovirus as a vector to induce production of SARS-CoV-2 S in the body ^11–15^. The Novavax vaccine (Nuvaxovid), a protein subunit vaccine, uses recombinant S proteins attached to a lipidic matrix for immunisation ^16–18^. These vaccines have been shown to be effective in preventing severe COVID-19, with the mRNA vaccines exhibiting highest efficacy rates (∼95%) albeit relatively short-lived ^1, 19^.

All currently licensed COVID-19 vaccines require refrigeration to maintain their stability and potency and depend on a functioning cold-chain ^20, 21^. This renders distribution and storage logistics of the vaccines challenging, especially in areas with limited access to refrigeration, which includes most developing nations with often remote or impoverished regions. In fact, cold-chain logistics issues are one of the main sources for vaccine spoilage and wastage ^22–24^. Therefore, developing thermostable vaccines that are not dependent on cold-chains, would greatly simplify the vaccine roll-out process and increase accessibility to vaccines globally.

Nanoparticle-based vaccines hold great promise for overcoming the limitations of current vaccine technologies ^25–27^. Shortly before the pandemic, we introduced ADDomer, a synthetic self-assembling protein nanoparticle platform for highly efficient vaccination by genetically encoded multiepitope display ^28^. ADDomer derives from a single protein component of adenovirus, which forms pentons at the vertices of the viral capsid, providing a base for the attachment of the adenoviral fibre ^29^. When produced in isolation, 60 copies of this penton-base protomer spontaneously self-assemble into a dodecahedron comprising 12 pentons. This adenovirus derived dodecamer, or ADDomer, is stable at temperatures exceeding 45°C and can be stored at ambient temperature for prolonged periods ^28^. We showed that exposed loops on the ADDomer surface can function as insertion sites for rationally selected immunogenic peptide epitopes that range in sequence and size, resulting in potent vaccine candidates against a range of human and veterinary infectious diseases^28, 30^.

In this study, initiated during the pandemic lockdowns, we set out to develop ADDoCoV, a thermostable ADDomer-based COVID-19 vaccine, with the potential to overcome limitations associated with the cold-chain challenge, while maintaining the advantages of ADDomer notably the ease of production of a one-component particle harbouring the genetically encoded antigens. We validate our ADDoCoV design by near-atomic resolution electron cryo-microscopy (cryo-EM) and molecular dynamics (MD) in a hybrid approach. We confirm authenticity and accessibility of the immunogenic epitope displayed by using Ribosome Display ^31^, a powerful *in vitro* selection technique, to generate neutralising antibodies *in vitro* from a naïve nanobody library. We demonstrate the prowess of our ADDoCoV design by immunising mice, eliciting antigen-specific IgA and IgG antibody responses *in vivo,* notably also by intranasal vaccination, a route unparalleled in terms of ease of administration, and with the potential to induce stronger and more long lasting indirect effects, especially in the context of a largely immune-primed global human population. Taken together, our approach provides a blueprint for the design of thermostable, cost-effective, convenient to manufacture, easy to administer, single-component vaccines, with the potential to prevent and combat infectious disease outbreaks also in resource-limited settings.

## Results

### Self-assembling thermostable ADDomer-based COVID-19 candidate vaccine ADDoCoV

The SARS-CoV-2 virion surface is decorated by S, a trimeric glycoprotein mediating cell attachment and infection ^32–34^. Each S monomer contains a receptor binding domain (RBD) comprising the receptor binding motif (RBM). In the open form adopted by S, the RBM is positioned to interact tightly with the cellular receptor, angiotensin converting enzyme 2 (ACE2) (Fig. 1a). Early in the pandemic when the sequence of SARS-CoV-2 S became available, we used alignments with SARS-CoV S to delineate peptide regions in the RBM which we could use as a putative antigenic epitope for genetic insertion into the protomer we had designed that forms the ADDomer nanoparticle ^28^. An epitope (AH) encompassing 33 amino acids of the SARS-CoV-2 Wuhan RBM spanning residues Y505 to Y473 inclusive (Fig. 1) was introduced into the insertion site of the variable loop (VL) of the protomer (Fig. 1b). The protomer adopts a bipartite structure (Fig. 1b,c), made up of a crown domain containing flexible loops and a jellyfold domain mediating multimerisation into pentons as well as the formation of the dodecahedron by establishing inter-penton contacts. Modelling by Rosetta design and molecular dynamics (MD) simulations revealed that AH exhibits considerable conformational flexibility that likely promotes antibody binding (Fig. 1c).

**Fig. 1:**
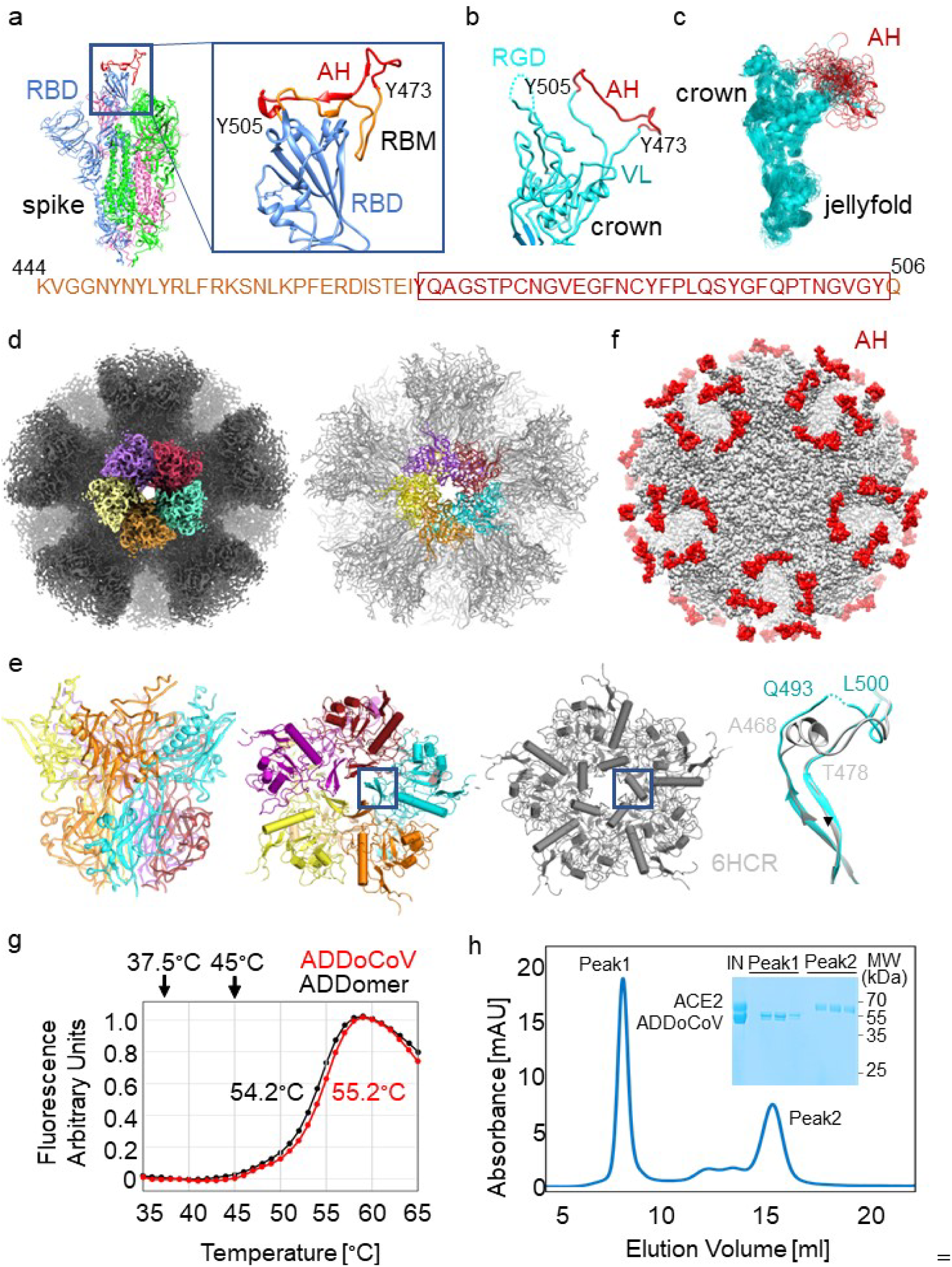
Self-assembling thermostable ADDoCoV candidate vaccine. **(a)** SARS-CoV-2 S protein (magenta, green and blue) shown in the open conformation (PDBID 7A94) ^89^. The zoom-in (right) on the RBD in the ‘up’ position (blue) depicts an ordered RBM (AA sequence provided below). The AH epitope (residue 473-505) is highlighted in red. **(b)** The ADDomer protomer crown domain (cyan) is shown. VL, variable loop; RGD, arginine-glycine-aspartate loop. AH (red) was inserted in VL. **(c)** MD simulations of the ADDoCoV protomer showing highly defined crown domain (cyan) while the AH epitope (red) in VL samples a range of conformations (RGD loop omitted for clarity). **(d)** Cryo-EM density (left) and model (right) of ADDoCoV. Five protomer (purple, firebrick, cyan, orange, yellow) form one penton. Twelve pentons form the nanoparticle. **(e)** One penton is depicted in a side view (left) and from the top (middle), coloured as in panel d. For comparison, a penton from the ADDomer scaffold (PDBID 6HCR)^28^ is shown (grey). A central region is boxed. Overlay of the boxed regions highlights unfolding of a central α-helix in the ADDoCoV protomer crown domain. **(f)** Model of ADDoCoV (grey) with AH epitopes coloured in red. **(g)** Thermal unfolding curves of ADDomer scaffold (black) and ADDoCoV (red) demonstrate high thermotolerance. Melting temperatures are indicated. **(h)** SEC profile and Coomassie-stained SDS PAGE section (inset) showing ADDoCoV (Peak1) and ACE2 (Peak2) eluting separately. IN, injected sample.

AH containing protomer was produced following our established protocol ^30^ resulting in highly purified ADDoCoV adopting the dodecahedral structure characteristic of this protein nanoparticle (Supplementary Fig. 1). We determined the cryo-EM structure of ADDoCoV at 2.36 Å resolution, providing near-atomic insights (Fig. 1d, Supplementary Figs. 2,3, Supplementary Table 1, Supplementary Movie 1). In a previous X-ray crystallographic study, a central α-helix had been identified, thought to stabilize the adenoviral penton by coordinating a bivalent ion, Ca^2+^, via glutamate residues ^35^. In ADDoCoV, this α-helix seemingly underwent a helix-to-disorder transition and cation coordination was not observed (Fig. 1e). ADDoCoV contains 60 AH epitopes exposed on the nanoparticle surface in flexible loops, available for antibody binding (Fig. 1f). We probed the dynamics of ADDoCoV by MD simulations guided by the cryo-EM structure. For about a third of the simulated trajectory, the AH epitope adopted a conformation closely resembling the arrangement observed in the open form of SARS-CoV-2 S, with the RBD in the ‘up’ position, positioned to engage ACE2 (Fig. 1e, Supplementary Fig. 4).

**Fig. 2:**
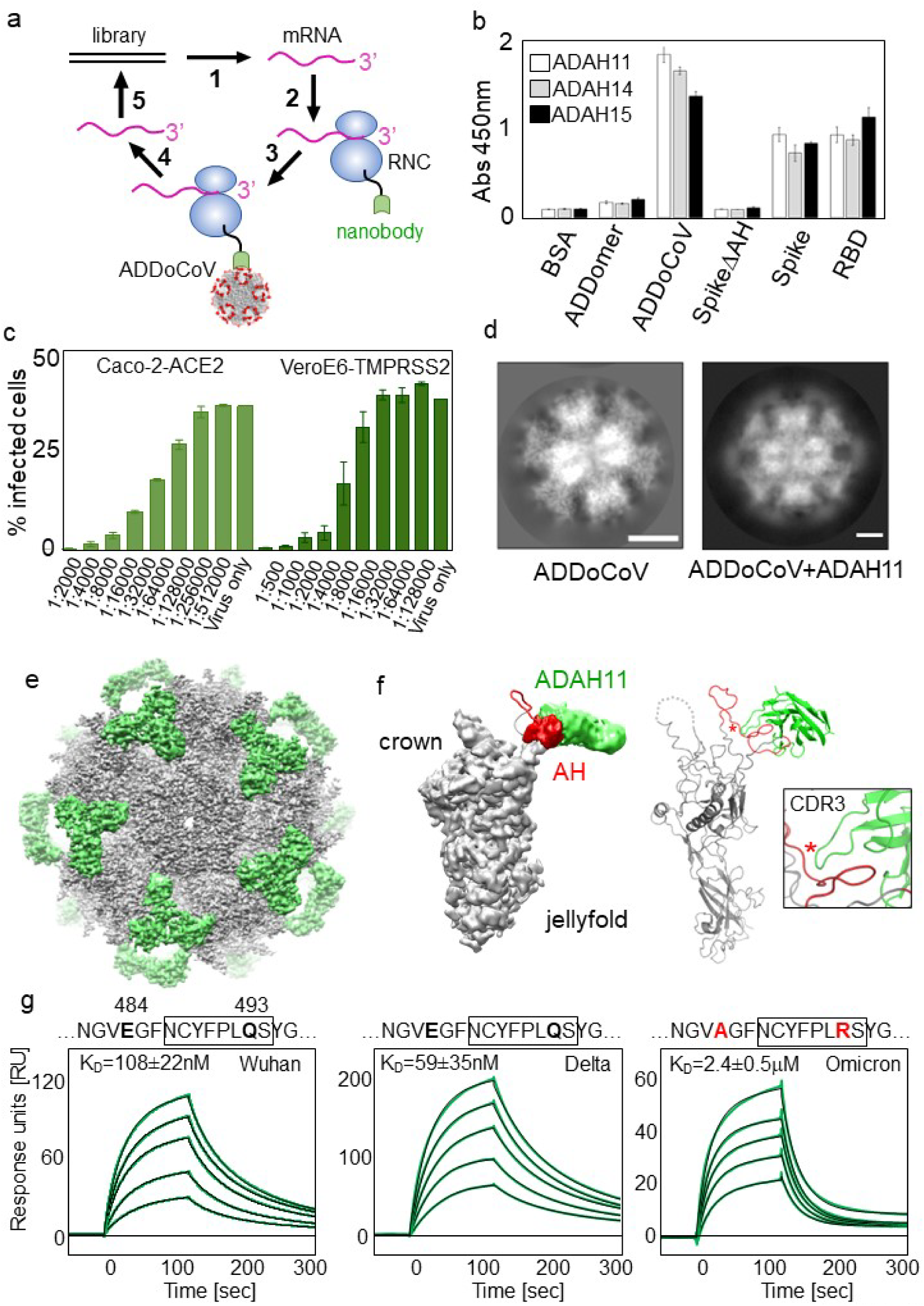
*In vitro* generated neutralising nanobodies validate ADDoCoV design. **(a)** Ribosome display in a schematic view. A DNA library encoding nanobodies is transcribed (1) and translated (2) *in vitro*. The stop codons in mRNAs are deleted. Resulting ribosome nascent-chain complexes (RNCs) displaying nanobodies are used for panning (3) against ADDoCoV antigen. After washing, the mRNA of bound RNCs is eluted (4) and DNA recovered by RT-PCR (5). The process is iterative. **(b)** ELISA of three selected nanobodies evidencing binding to ADDoCoV, S and RBD, but not to BSA, ADDomer scaffold, or a S mutant with AH deleted (SpikeΔAH). **(c)** SARS-CoV-2 neutralisation by ADAH11 using ACE2 expressing Caco-2 (left) and TMPRSS2-expressing Vero E6 cells (right). Dilutions indicated were sampled in triplicates. **(d)** Reference-free 2D class averages of ADDoCoV (left) and of ADDoCoV-ADAH11 nanobody complex (right) evidence halo of density corresponding to bound nanobody. **(e)** Cryo-EM structure of ADDoCoV (grey density) with bound nanobody (green). **(f)** Symmetry expansion (left) of ADDoCoV protomer (grey) with nanobody (green, filtered to ∼10Å resolution) bound to AH epitope (red). The corresponding molecular model (right) suggests ADAH11 recognising a central section of AH. Location of an arginine residue in the ADAH11 CDR3 is marked (asterisk). **(g)** SPR of ADAH11 binding to immobilised Wuhan (left), Delta (middle) and Omicron (right) RBDs, at concentrations 50 nM to 250 nM for Wuhan and Delta, and 1 uM to 3 uM for Omicron. Epitope sequences are provided (top). Section bound by ADAH11 is boxed. Mutations in the Omicron RDB are highlighted (red). K_D_s are indicated.

**Fig. 3:**
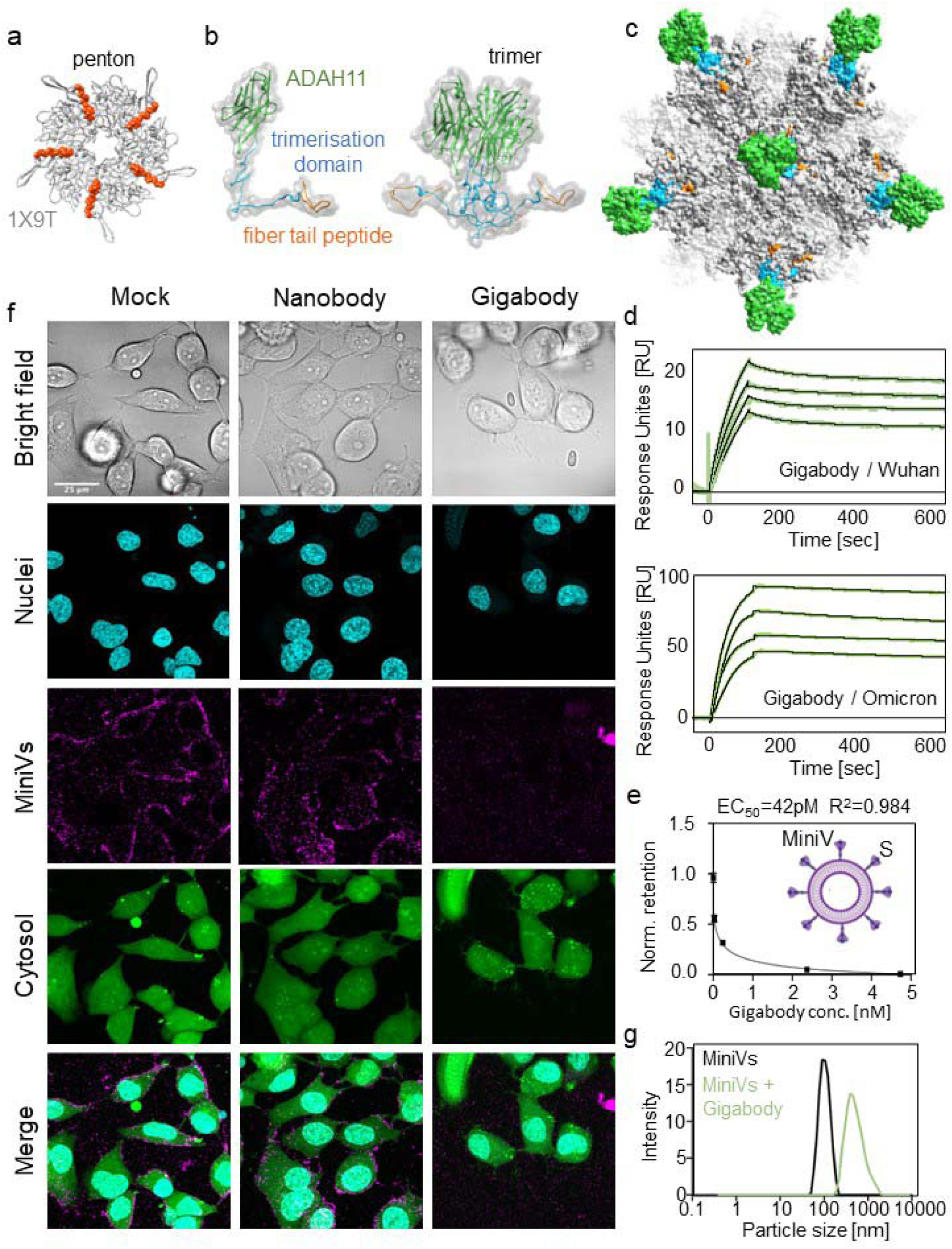
Multivalent picomolar affinity Gigabody against SARS-CoV-2 RBM. **(a)** Top view of an adenovirus penton base (grey) in complex with N-terminal fibre peptide (orange) is shown (PDBID 1X9T)^35^. **(b)** Protein engineering of ADAH11 nanobody (green) with a C-terminal T4 foldon trimerisation domain (blue) and fibre tail peptide (orange) results in a trimeric complex (right). **(c)** Gigabody comprising 12 trimers bound via fibre tail peptides to pentons, displaying a total of 36 ADAH11 nanobodies. **(d)** SPR of Gigabody interactions with immobilised Wuhan (above) and Omicron (below) RBDs evidence very slow dissociation, consistent with tight (picomolar) binding. Gigabody concentrations of 0.5 to 2.0 nM (with immobilised Wuhan RBD) and of 1 to 2.5 nM (with immobilised Omicron RBD) were used. **(e)** Quantification of MiniV retention in ACE2-expressing A549 cell monolayers 2.5 hours after incubation. Competitive binding of Gigabody to MiniV-presented S was assessed in a serial dilution series. Graph shows mean standard deviations from three technical replicates. IC50 is indicated. **(f)** Laser scanning confocal microscopy images of ACE2-expressing A549 cells 2.5 hours after incubation with synthetic SARS-CoV-2 MiniVs decorated with S glycoprotein. MiniVs were either left untreated or exposed to 500 nM ADAH11 nanobodies or 1.6 nM Gigabody (corresponding to equal final protein concentration) for 30 min before addition to the cell cultures. Scale bar is 50 µm. **(g)** Dynamic light scattering analysis of Gigabody-mediated MiniV aggregation. MiniVs hydrodynamic size distribution is shown for untreated controls, and MiniVs that were pre-treated with 1.6 nM Gigabody for 30 min, respectively.

A key feature of the self-assembling ADDomer scaffold resides in its thermostability ^28^. We observed virtually identical melting temperatures of ADDomer and ADDoCoV, confirming that, irrespective of AH epitope insertion, thermostability is maintained (Fig. 1g). The AH epitopes displayed on the ADDoCoV nanoparticle comprise 33 amino acid residues of the SARS-CoV-2 RBM which is itself about 60 residues long. We deliberately chose the shorter AH epitope for ADDoCoV to preclude potentially detrimental effects that could be caused by ADDoCoV sticking to cellular receptor ACE2. Size exclusion chromatography (SEC) of a mixture of ADDoCoV and highly purified, recombinant ACE2 showed no association with ACE2, notwithstanding the presence of 60 copies of the AH epitope on the ADDoCoV nanoparticle (Fig. 1h).

In summary, we designed ADDoCoV comprising 60 copies of AH, a SARS-CoV-2 RBM derived epitope, determined ADDoCoV architecture and integrity at near atomic resolution, sampled the dynamics of the AH epitopes displayed on the ADDoCoV surface and demonstrated thermostability of this COVID-19 nanoparticle vaccine candidate.

### *In vitro* generated SARS-CoV-2 neutralising nanobody binders by Ribosome Display

The rationale for the ADDoCoV vaccine design is to elicit antibodies that can bind the RBM, and thus prevent SARS-CoV-2 attachment to ACE2, neutralising the virus. A prerequisite for this is authenticity and accessibility of the AH epitope in the context of the ADDoCoV nanoparticle. To validate our design, we used Ribosome Display to select antibody binders from a naïve synthetic nanobody library, with ADDoCoV as an antigen (Fig. 2a). In Ribosome Display, a DNA library encoding for nanobodies is transcribed and translated *in vitro* ^31^. In the library, the stop codons are deleted and replaced with a DNA sequence encoding an oligopeptide spacer. Thus, *in vitro* transcription and translation gives rise to ribosome nascent chain complexes (RNCs) coupling the genotype (mRNA) to the phenotype (nanobody), tethered to the ribosome. RNCs comprising specific nanobody binders are rapidly selected by panning on ADDoCoV immobilised on a surface. After washing away unbound RNCs, the remaining mRNA is recovered by dissociating the bound RNCs. Reverse transcription and PCR regenerates a DNA pool enriched for specific nanobody binders (Fig. 2a). By ELISA, we identified nanobodies that bound ADDoCoV as well as SARS-CoV-2 S and RBD, but not bovine serum albumin (BSA), native ADDomer scaffold, and S lacking the AH epitope (Fig. 2b, Supplementary Table 2). A nanobody identified in this way, ADAH11, showed efficient virus neutralisation in live SARS-CoV-2 infection assays using two different ACE2-expressing cell lines (Caco-2-ACE2 and VeroE6-TMPRSS2) (Fig. 2c).

ADAH11 was expressed and purified to homogeneity and tested for binding to highly purified ADDoCoV by SEC confirming complex formation (Supplementary Fig. 5). Purified ADDoCoV-ADAH11 complex was analyzed by cryo-EM (Fig. 2d-f, Supplementary Fig. 6, Supplementary Table 3). Comparison of reference-free 2D class averages of ADDoCoV-ADAH11 or ADDoCoV, respectively, clearly indicated additional density for the nanobody containing complex (Fig. 2d). The cryo-EM structure of the ADDoCoV-ADAH11 complex revealed nanobody binding to the crown domains comprising the AH epitope (Fig. 2e,f, Supplementary Fig. 6e). The arrangement of the pentons within the dodecahedron locates the AH epitopes in apparent triangles on the ADDoCoV surface. This is reflected by the triangular shape of the extra density stemming from nanobody binding (Fig. 2e). The limited resolution of the cryo-EM density in these more flexible outer regions only allowed rigid body docking of the nanobody, suggesting that ADAH11 is interacting with a central segment of the AH epitope in the protomer crown domain (Fig. 2f). Nanobody binding to cognate antigen is typically dictated by complementarity-determining region 3 (CDR3). In ADAH11, CDR3 comprises an arginine residue, R105 (Fig. 2f, Supplementary Table 2). By using surface plasmon resonance (SPR) by Biacore, we characterised ADAH11 binding to the SARS-CoV-2 S RBDs of the original Wuhan virus, as well as to the currently dominating Variants of Concern (VOCs) Delta and Omicron (Fig. 2g). ADAH11 bound immobilised S RBD of Wuhan and Delta, both, with similar, low nanomolar affinity (K_D_ of 108 nM and 59 nM, respectively). Omicron S RBD, in contrast, was bound significantly less tightly (K_D_ of 2.4 μM). Of note, ADAH11 binding to Wuhan and Delta S would juxtapose R105 in CDR3 with a glutamine residue in the RBMs. In Omicron, in contrast, this glutamine is mutated to arginine (Fig. 2g), The resulting juxtaposition of two positively charged arginine residues in RBM and CDR3 could thus contribute to significantly reduced binding of Omicron S RBD by ADAH11.

To summarise, a nanobody specific for an epitope derived from the RBM in the SARS-CoV-2 RBD was selected by Ribosome Display. This *in vitro* generated nanobody bound ADDoCoV, cross-reacted with the RBM in SARS-CoV-2 S, and neutralised live SARS-CoV-2 in cell-based infection assays, validating the authenticity and accessibility of the RBM-derived AH epitope displayed on the ADDoCoV nanoparticle vaccine.

### ADDomer-based ultrahigh-affinity Gigabody displaying SARS-CoV-2 nanobody binders

In adenovirus, the penton represents the base for attachment of the adenoviral fibre proteins that form characteristic protrusions at the vertices of the adenoviral capsid. The fibre adopts a trimer of three identical fibre proteins. Attachment to the penton base is mediated by a highly conserved, proline- and tyrosine-rich N-terminal fibre tail peptide present on each of the monomers (Supplementary Table 4). The fibre tail peptide binds to a tailormade fibre tail peptide-binding cleft on the penton base. In a previously reported crystal structure of an adenoviral penton bound to isolated fibre tail oligopeptides, all five binding clefts were occupied (Fig. 3a, Supplementary Fig. 8) ^35^. In the adenovirus, binding of the trimeric fibre to the penton base will result in two of five clefts remaining unoccupied.

The nanobodies we selected by Ribosome Display neutralised live SARS-CoV-2, most likely by blocking interactions with the ACE2 receptor due to steric hindrance. The nanobodies we obtained in this way were characterised by binding affinities of about 100 nM to their target antigen (Fig. 2g, Supplementary Fig. 7). Multimerisation of a nanobody can result in much tighter binding by increased avidity. We set out to exploit the principles of adenoviral fibre attachment using nanobody ADAH11 as a starting point, with the aim of creating an ultra-high affinity superbinder, ‘Gigabody’, displaying multiple copies of ADAH11, with the potential to forestall SARS-CoV-2 infection, which conceivably, could be utilised for passive immunisation.

We had designed ADDoCoV based on a penton base protomer derived from human adenovirus serotype Ad3, which can efficiently self-assemble into a dodecahedron ^28^. For Gigabody, we chose a different protomer, derived from chimpanzee adenovirus AdY25 to form the ADDomer (Supplementary Table 5). Using the adenovirus fibre as a blueprint, we designed a nanobody trimer by fusing a T4 phage derived trimerisation domain (T4 foldon) preceded by the AdY25 fibre tail, to the N-terminus of ADAH11 (Fig. 3b, Supplementary Table 6). Next, Gigabody was produced by mixing the trimers with AdY25-derived ADDomer, purified by SEC and dodecahedron formation confirmed by negative-stain EM (Supplementary Fig. 8). Due to the 3:5 symmetry mismatch of ADAH11 trimer and penton, the trimer structure cannot be resolved at high resolution by cryo-EM, and computational modeling was used to illustrate the geometry of the Gigabody nanoparticles (Fig. 3c, Supplementary Fig. 9, Supplementary Movie 2). Fully occupied Gigabody presents 36 ADAH11 nanobodies arranged in 12 trimers, which should substantially increase binding to the cognate AH epitope by increasing avidity. We tested Gigabody binding to SARS-CoV-2 Wuhan S RBD by SPR. As expected, binding improved substantially, from about 100 nM for momomeric ADAH11 to picomolar for the Gigabody, driven by very slow dissociation (Fig. 3d). Moreover, with monomeric ADAH11, we had observed a significant drop in binding affinity from Wuhan S RBD (100 nM) to Omicron S RBD (2.4 μM) whereas Gigabody binding to Wuhan and Omicron was virtually identical, in the picomolar range (Fig. 3d). This indicates that Gigabody, by presenting multiple copies of ADAH11, can rescue the comparatively low, micromolar affinity binding by the nanobody to Omicron S RBD (Fig. 3d).

We tested the capacity of Gigabody to abrogate virion attachment to ACE2 expressing cells. We used synthetic minimal SARS-CoV-2 virions decorated with highly purified S glycoproteins (SARS-CoV-2 MiniVs) as a model system, affording complete control of experimental parameters ^36^. We had used synthetic MiniVs previously to reveal fatty-acid coupled adaptive immunogenicity of SARS-CoV-2 ^36^. SARS-CoV-2 MiniVs faithfully recapitulate viral attachment and can be studied in a regular laboratory setting (biosafety level 1), in contrast to live virus. We assessed competitive binding of Gigabody to MiniV-presented S in a serial dilution (Fig. 3e) and analysed attachment of SARS-CoV-2 MiniVs to ACE2-expressing A549 cells exposed to Gigabody by laser scanning confocal microscopy (Fig. 3f). We observed quantitative inhibition of SARS-CoV-2 MiniV cell attachment at a Gigabody concentration of 1.6 nM (Fig. 3f). Previously, we had determined EC_50_ of ADAH11 nanobody in terms of MiniV retention as 117 nM ^36^. Gigabody EC_50_ is 42 pM (Fig. 1e), which is 300-fold lower, closely mirroring the respective binding properties of single nanobody and Gigabody, respectively, in SPR measurements (Fig. 2g, Fig 3d). Due to the presence of multiple nanobody trimers bound to the pentons, Gigabody could in theory induce virion aggregation, similar to agglutination. We analysed the hydrodynamic size distribution of SARS-CoV-2 MiniVs by dynamic light scattering, confirming aggregation following Gigabody addition, with particle sizes increased to ∼1000 nm from the diameter of a single virion (∼100 nm) (Fig. 3g).

Taken together, we have mimicked the design of the adenoviral fibre, and its attachment mechanism in the adenovirus, to generate a Gigabody nanoparticle decorated with multiple copies of trimerised ADAH11. By avidity, Gigabody binds the cognate target in the RBM of SARS-CoV-2 S with greatly enhanced, picomolar affinity as compared to nanomolar binding by ADAH11 nanobody alone. Moreover, Gigabody binding to the RBMs in Wuhan S and Omicron S is virtually identical, while ADAH11 alone binds Omicron with significantly reduced affinity as compared to Wuhan, presumably due to the mutations accrued by Omicron in the RBM. Finally, Gigabody effectively abolishes attachment of SARS-CoV-2 MiniVs in cell-based assays and can mediate virion agglutination. Intriguingly, Gigabody thus may represent an attractive avenue for passive immunisation, based on the same nanoparticle scaffold concept, ADDomer, that we used for ADDoCoV, exploiting assembly principles of the adenovirus from which ADDomer is derived, and utilising antibody binders generated *in vitro* against ADDoCoV used as an antigen.

### ADDoCoV immunisation experiments *in vivo* in mice

Traditional routes of vaccine administration, such as intramuscular (IM), subcutaneous (SC) and intradermal vaccination, generally induce significant concentrations of antigen-specific IgG that are detectable in the recipient’s serum. In the current study, we compared traditional IM or SC vaccination with intranasal (IN) vaccination as this route may also induce strong systemic responses as well as significant levels of antigen-specific antibody detectable in mucosal secretions. We hypothesised that IN vaccination might be beneficial for a SARS-CoV-2 vaccine, as it could increase front-line mucosal defences against a pathogen that infects the respiratory tract and thus impact on viral transmission more effectively than the vaccines currently in use. Therefore, we tested the immunogenicity of ADDoCoV using a homologous prime-boost protocol (Fig. 4a). Specifically, we tested the immunogenicity of ADDoCoV in mice as compared to the naïve ADDomer nanoparticle as a control, when administered via SC, IM and IN routes. As shown (Fig. 4b-d), after the vaccine prime, the ADDoCoV formulation resulted in 100% seroconversion (*n* = 10 mice/group) regardless of the route of administration. Elevated concentrations of anti-RBD specific serum immunoglobulin-G (IgG) were detected during study weeks 3, 6 and 9. When compared to baseline, anti-RBD IgG antibody titres in serum in week 9 after vaccination were significantly increased across all conditions (\*\*\*\**p*≤0.0005) (Fig. 4e).

**Fig. 4:**
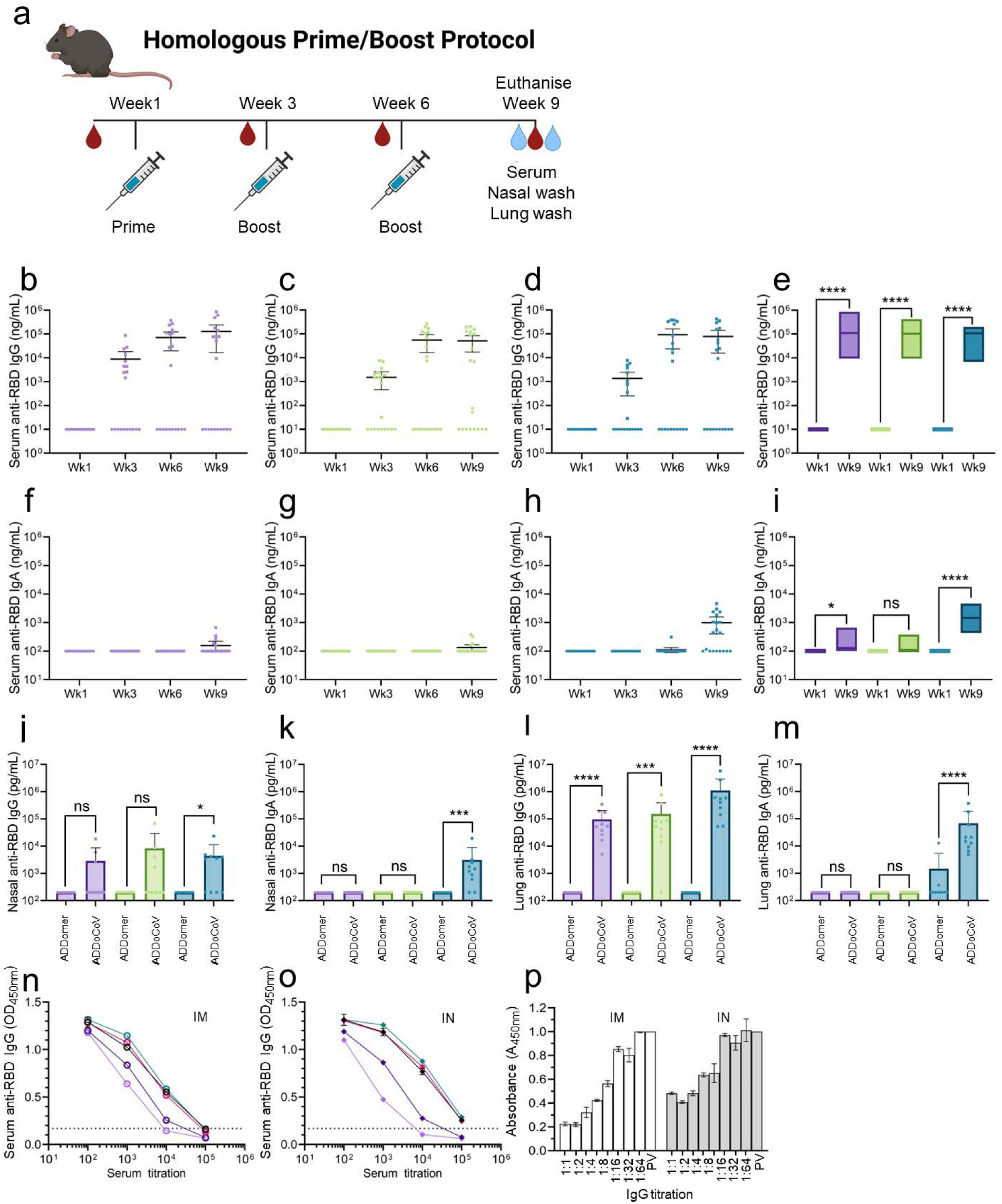
ADDoCoV *in vivo* immunisation elucidates nasal vaccination route. **(a)** Schematic of immunisation schedule and end point. **(b–e)** Determination of anti-RBD specific IgG binding antibodies induced through sub-cutaneous immunisation (purple), intra-muscular immunisation (green) and intra-nasal immunisation (blue). **(f–i)** Comparison of anti-RBD specific IgA binding antibodies induced through sub-cutaneous (SC) immunisation, intra-muscular (IM) immunisation and intra-nasal (IN) immunisation. **(j,k)** Determination of anti-RBD specific IgG binding antibodies in nasal washes induced by IM or IN immunisation. **(l,m)** Determination of anti-RBD specific IgG binding antibodies in lung washes induced by IM or IN immunisation. Control unvaccinated animals are represented by •, vaccinated animals by ▪ symbols. Statistical analysis utilised two-sided Mann–Whitney tests. **(n,o)**: Assessment of cross-variant binding for total IgG antibodies induced through IM (n) or IN (o) immunisation. Black (Alpha variant), pink (Delta), teal (Wuhan), purple (Beta), mauve (Omicron). Dotted line represents upper 99% CI of blank controls. **(p)** Surrogate virus neutralisation assay (sVNT) ^38^ assessing total IgG antibodies induced through IM (white bars) or IN (grey bars) immunisation, normalised to pre-vaccination sample (PV) included as control (bars far right). Dilutions indicated were sampled in triplicates.

Given the important role that immunoglobulin A (IgA) plays in defence against mucosal pathogens ^37^, we also measured the serum anti-RBD IgA response after ADDomer and ADDoCoV vaccination. Detectable anti-RBD IgA was also induced in serum (Fig. 4f-h), however the IgA response developed more slowly than the IgG response, with anti-RBD IgA antibody only being significantly elevated at week 9 in the SC and IN groups compared to the week 1 baseline control (Fig. 4i). Importantly, only the IN group showed 100% seroconversion after ADDoCoV administration, with the IN routes also resulting in the highest anti-RBD IgA response in serum. Taken together, these data show that the ADDoCoV vaccine is immunogenic and elicits antigen-specific antibody responses in the serum of vaccine recipients, irrespective of the route of administration. However, the magnitude, kinetics and isotype of the elicited antibody response are influenced by the route of administration.

To gain a more in-depth understanding of the elicited antibody response after SC, IM and IN vaccination with ADDoCoV at the end of the study, we collected nasal (Fig. 4j,k) and lung washes (Fig. 4l,m) to quantify the anti-RBD IgG and IgA responses in mucosal secretions. As expected, systemic routes of vaccination failed to induce detectable anti-RBD IgG and IgA responses in nasal secretions with only the IN administration of ADDoCoV resulting in significantly elevated concentrations of IgG (\**p*≤0.05) and IgA (\*\*\**p*≤0.0005) compared to the ADDomer control vaccine. Interestingly, a distinct pattern of anti-RBD antibody induction was observed in lung washes. Here we found that significant levels of anti-RBD IgG was induced regardless of the route of vaccine administration. However, despite the IM, SC and IN groups all having significant levels of anti-RBD in lung washes, it should be noted that the IN group had the highest. Interestingly, only in the IN group were significant levels of anti-RBD IgA induced in lung washes (Fig. 4m). Taken together these results suggest that IN immunisation may induce mucosal antibody responses more efficiently than systemic routes of administration and support the use of this route of administration to enhance interruption of acquisition and onward transmission of infection.

Next, we sought to evaluate whether the antibodies induced by ADDoCoV vaccination were cross-reactive against a range of clinically relevant SARS-CoV-2 variants (Wuhan, Alpha, Beta, Delta, and Omicron) which would be important for broad protection, given the rapid and continuing evolution and diversification of SARS-CoV-2 in the human population. IgG antibodies induced following both IM and IN administration of the vaccine bound all SARS-CoV-2 S RBDs tested, with close to identical binding observed to Wuhan, Alpha and Delta RBDs, and reduced binding observed for Beta and Omicron RBDs (Fig. 4n,o). Of note, a reduction in binding to Omicron RBD, as compared to Wuhan and Delta RBDs, was also observed for ADAH11, the nanobody generated by Ribosome Display (Fig. 2g), suggesting that our *in vitro* selection may reproduce some of the antibody binding characteristics likewise occurring *in vivo*.

We showed that the ADDoCoV vaccine is immunogenic and induces antibodies in serum and mucosal secretions which are cross reactive against diverse SARS-CoV-2 S protein variant RBDs. Next, we used serum IgGs to perform a surrogate virus neutralisation assay (sVNT) ^38^, based on competition with Wuhan RBD bound to immobilised ACE2. We observed moderate neutralising activity in our tests (Fig. 4p). Antibody-mediated neutralisation is thought to be important in protection against infection by SARS-CoV-2 ^39^. At the same time, absence of strong neutralisation does not equate lack of protection by serum antibodies, as even entirely non-neutralising antibodies, which bind target proteins in pathogens specifically, can confer strong protection through antibody-dependent cellular functions ^40–42^. We note that our *in vitro* generated nanobody binder, ADAH11, showed strong viral neutralisation (Fig. 2c). The serum antibodies, whilstspecifically binding S RBM, may be characterised by moderate binding affinity, restricting their performance in the sVNT assay. It is conceivable that iterative adjustment and refinement of the vaccine design (Fig. 5), for instance by incorporating additional epitopes, could result in stronger binding and more efficient neutralisation by the antibodies generated.

**Fig 5:**
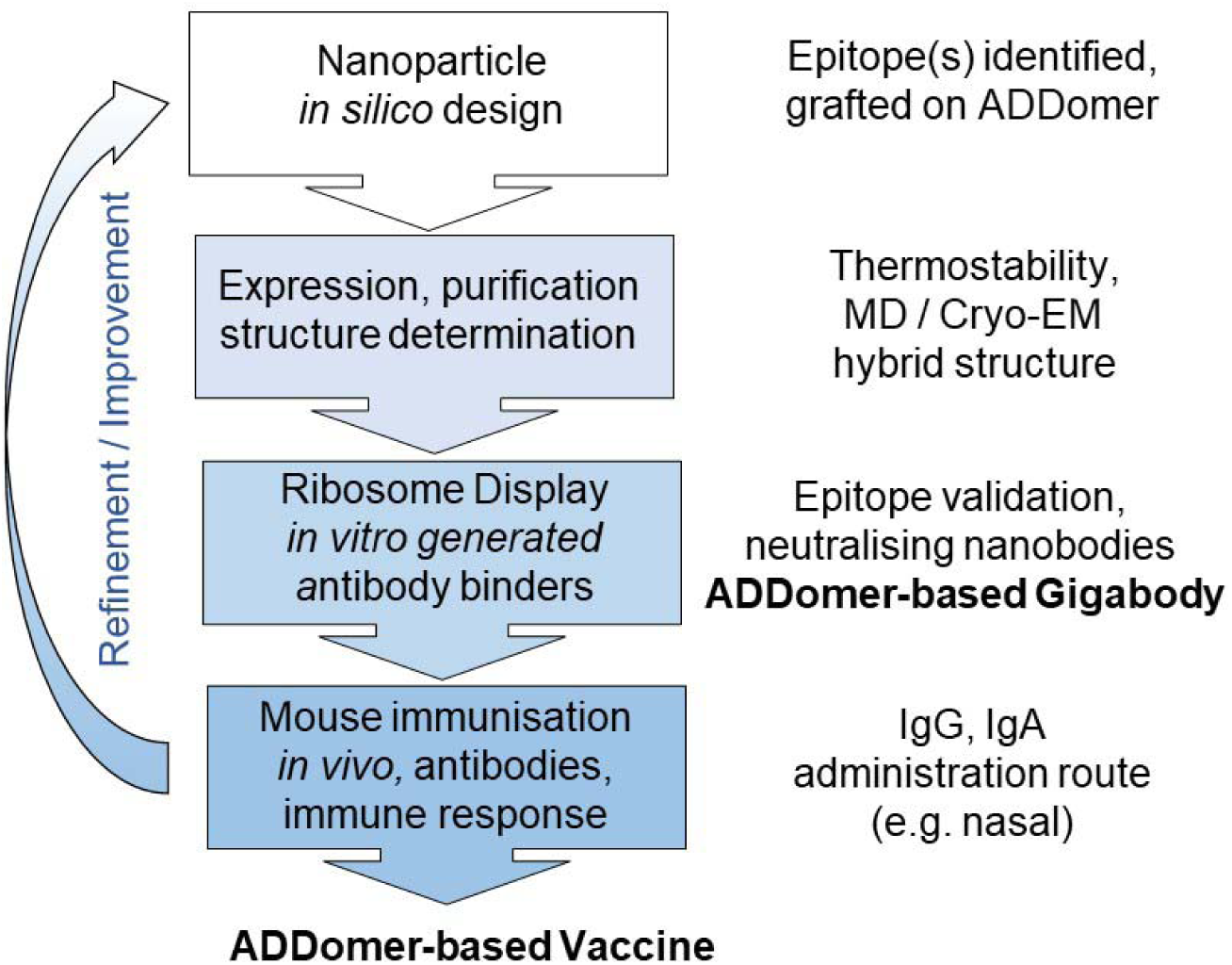
Pipeline for ADDomer-based vaccine and Gigabody design. Integrating *in silico* design, cryo-EM, MD simulations, *in vitro* selection by Ribosome Display, synthetic SARS-CoV-2 virions and live virus neutralisation, and validation in an animal model, for generating ADDomer-based nanoparticle therapeutics for active (vaccine) and passive (Gigabody) immunisation. With established protocols for each step, the process from immunogenic epitope identification and grafting onto ADDomer, until release to animal studies, is rapid and can be completed in about five weeks. The process can be repeated iteratively to refine and optimise, for instance by including diverse additional B and T epitopes in the design.

## Discussion

In summary, we developed ADDoCoV, a thermostable, self-assembling, self-adjuvanting multi-epitope display nanoparticle vaccine candidate against SARS-CoV-2, the causative agent of COVID-19 utilizing an efficient and rapid validation pipeline (Fig. 5). Our pipeline combines synthetic, structural and computational methods for epitope selection and vaccine candidate design with *in vitro* selection of neutralising nanobodies to confirm authenticity and accessibility of the epitope displayed, followed by *in vivo* immunisation studies in mice to characterise the immune response and how this is influenced by route of administration. We note that our *in vitro* selection by Ribosome Display generated nanobody binders that exhibited properties similar to the antibodies in the sera of immunised mice. This point is relevant in the context of the principle of 3Rs (reduce, replace, refine) to limit animal use for biomedical research, here for prescreening suitable vaccine candidates with the desired properties regarding the antigen displayed. Our mouse experiments elucidated nasal administration as a viable route for ADDoCoV vaccination, eliciting specific serum IgG and mucosal IgA responses *in vivo*. Moreover, making use of the neutralising nanobodies we selected *in vitro*, we created Gigabody, a novel ‘superbinder’ nanoparticle exploiting the same adenovirus-derived dodecamer scaffold concept that we used for ADDoCoV. Gigabody displays multiple trimers of a neutralising nanobody, effectively blocks virion attachment to ACE2 expressing cells *in vitro* and could potentially be administered similar to ADDoCoV via the nasal route, for passive immunisation. Because we used ADDomer-forming protomers of different origin (human Ad3 serotype and chimpanzee AdY25) as a scaffold for the ADDoCoV vaccine and Gigabody superbinder respectively, it is conceivable to deploy both vaccine and superbinder for active and passive immunisation in a given treatment regimen, circumventing potential pre-immunity issues. In this context it is noteworthy that several adenovirus serotypes have been identified comprising penton base proteins that can adopt dodecahedrons and could be used as scaffolds ^29^.

In our mouse immunisation experiments, administration of ADDoCoV was sufficient to induce serum anti-RBD IgG production by subcutaneous, intramuscular, and intranasal routes, demonstrating that ADDoCoV is capable of inducing systemic immune responses to the relevant target. Since the ADDomer has been shown to self-adjuvant ^28^, adjuvant was not included in our experiments. Serum anti-RBD IgG production was detectable at week 3 after a single dose and rose further following boosting at weeks 3 and 9. In all ADDoCoV treatment groups, significantly increased concentration of serum anti-RBD antibodies was detectable at week 9 compared to treatment-naïve animals at baseline.

Analysis of anti-RBD IgA responses at each time point revealed that protocol design, time and route of administration were important determinants of IgA induction. Serum anti-RBD IgA was detectable in peripheral blood samples only at week 9, following a complete prime-boost-boost protocol, showing a difference in seroconversion between IgG and IgA. Our results are consistent with maturation of the immune response following initial prime and subsequent boost treatments. Significant increases in serum anti-RBD IgA was observed only following subcutaneous or intranasal administration, while no significant increase in IgA production above baseline (treatment naïve) levels was detected in the intramuscular treatment group. This is interesting given that mucosal immunity is thought to be a critical aspect of SARS-CoV-2 infection ^43^, and supports what has been shown with other vaccines, that the recruitment of the appropriate physiological, e.g. mucosal, responses requires appropriate delivery of vaccine to relevant tissues.

The prevalence of antigen specific (anti-RBD) local/mucosal antibody production was assessed in nasal washes and bronchoalveolar lavage samples collected at week 9. Detection of vaccine induced local nasal IgG and IgA was limited to mice treated via intranasal ADDoCoV administration, whereas treatment via subcutaneous or intramuscular routes was inefficient in the generation of nasal mucosal antibody, as no increase in anti-RBD IgG or IgA was detectable above control levels. In contrast, lung mucosal IgG production was not dependent on the route of vaccine administration and was detected in ADDoCoV treated mice via all treatment routes. Induction of lung mucosal anti-RBD IgA production remained limited by the route of vaccine delivery, and was only detected in mice that received intranasal ADDoCoV administration. This is noteworthy as it further demonstrates specificity of induced mucosal responses according to route of administration, which was suggested by the presence of IgA in the serum of mice only after intranasal treatment. IgA is known to play a crucial role in the immune defense of mucosal surfaces, the first point of entry of SARS-CoV-2 ^44^.

Our current ADDoCoV vaccine design comprises 60 copies of the antigenic epitope AH, derived from the SARS-CoV-2 S RBM. The rationale for our design was to elicit an immune response that results in antibodies binding the RBM, sterically obstructing ACE2 binding. The SARS-CoV-2 neutralising nanobodies we selected *in vitro* by Ribosome Display compellingly validated our design, which was subsequently further underscored by the generation of anti-RBD antibodies, IgG and IgA, in vaccinated mice. Our synthetic ADDomer scaffold comprises three insertion sites per protomer ^28^, and only one is currently occupied by AH. Cellular immunity mediated by T cells is known to play an important role in the protection against viral infection, mediating effective viral clearance, elimination of virus-infected cells, and long-term disease protection. ADDomer was shown to drain to lymph nodes and is efficiently taken up by antigen presenting cells ^28^. Successful presentation of T cell epitopes, in addition to B cell epitopes, by ADDomer has been demonstrated recently for Type O foot-and-mouth disease virus, resulting in protective responses against the viral pathogen ^45^. A range of T cell epitopes in the SARS-CoV-2 proteome have been identified. It is likely that by expanding our ADDoCoV design to include validated SARS-CoV-2 T epitopes in the currently unoccupied insertion sites in the ADDomer scaffold, a T cell response can also be activated against SARS-CoV-2. Emerging SARS-CoV-2 VOCs, in particular Omicron, are characterised by multiple mutations, increasingly evading existing antibody responses, requiring updated versions of current vaccines to confer immunity. The ease of epitope insertion, and epitope alteration, on the genetic level renders the ADDomer scaffold particularly attractive for rapid, rolling update. We note that insertion of strings comprising several immunogenic epitopes in a row has been demonstrated for ADDomer ^28^. We propose rapidly updated ADDoCoV nanoparticles, comprising strings of the respective B and T epitopes, as attractive candidates for recurring vaccination against SARS-CoV-2 VOCs. Again, to avoid pre-immunity issues, scaffolds of different origin could be used for booster vaccinations.

Affordable production is a key prerequisite for broad vaccine distribution in resource-limited settings. Other nanoparticle-based SARS-CoV-2 vaccine candidates are often made up of one or several different S proteins, or their RBDs, which are coupled to nanoparticle scaffolds ^46–48^. For instance, a SARS-CoV-2 mosaic vaccine comprised 8 different S RBDs coupled to a separate nanoparticle ^46^. A different nanoparticle vaccine candidate displays S on a nanoparticle scaffold which itself is made up of different components, each produced in a different heterologous system ^47^. These nanoparticle vaccines share in common that all components, S, RBD and nanoparticles, need to be produced and purified separately, then combined and repurified, multiplying manufacturing runs and associated costs. In contrast, ADDoCoV relies on genetically encoded multiepitope display by a single, one-component particle, requiring one production run only using established manufacturing technology, significantly reducing costs and maintaining thermostability of the particle.

We utilised our pipeline to address SARS-CoV-2, but our approach can be applied to any other infectious disease-causing pathogen for which immunogenic epitopes are known to exist. We anticipate thermostable and affordable ADDomer-based nanoparticle therapeutics, both, vaccines for active, and Gigabodies for passive immunisation, designed, produced and validated as described here for ADDoCoV, to tackle many human and animal infectious diseases including pandemic outbreaks, present and future.

## Materials and Methods

### Protein production

#### ADDoCoV preparation

ADDoCoV was designed during the early pandemic before SARS-CoV-2 S structures became available, based on sequence comparison of the RBDs of SARS-CoV S, MERS-CoV S and SARS-CoV-2 S, and the structure of SARS-CoV bound to ACE2 receptor or neutralising antibodies, respectively ^49–52^. Variations of the oligopeptide sequence corresponding to the ACE2 receptor binding motif (RBM) were then inserted into the ADDomer scaffold as described previously^28^ and expressions carried out using the MultiBac baculovirus expression system following established protocols ^53–55^. ADDoCoV, the candidate here described comprises a 33 amino acid sequence (AH epitope) in the VL insertion site (Fig. 1a, Supplementary Table 1).

ADDoCoV purification was adapted a from a previously established protocol ^56^ Briefly pellets were resuspended in Resuspension Buffer (50 mM Tris pH 7.5, 150 mM NaCl, 2 mM MgCl_2_ Buffer, 1 ml per 2.5×10^7^ cells) supplemented by EDTA-free complete protease inhibitor (Roche). Lysate was prepared by three cycles of freeze-thawing, cleared by centrifugation (40,000g, 30 min), supplemented with Benzonase (Sigma-Aldrich) and incubated on ice for 2 hours. Precipitate was removed by centrifugation (4000g, 15 min), the supernatant passed through a 0.45 µm filter and subjected to size exclusion chromatography (SEC) using a XK 26/100 column (GE Healthcare). Fractions containing ADDoCoV were pooled and further purified by ion exchange chromatography (IEX) using a 5 ml Bio-Scale Mini Macro Prep High Q (Bio-Rad) equilibrated in Buffer A (50 mM Tris pH 7.5, 150 mM NaCl) and a linear salt gradient from 0.15 M to 1 M NaCl. Highly purified ADDoCoV eluted at ∼300 mM to 400 mM NaCl. Fractions were pooled and stored at ambient temperature or refrigerated (4°C). For animal studies, ADDoCoV particles were filtered through a 0.22 µm filter and further purified utilising Detoxi-Gel (ThermoFischer Scientific) to remove endotoxins and dialysed against PBS.

#### Receptor-Binding Domains (RBDs)

Biotinylated SARS-CoV-2 RBDs of Wuhan, Alpha, Beta, Delta and Omicron were expressed and purified as described ^57^.

#### ADAH11 expression and purification

A synthetic gene (Genscript) encoding for ADAH11 was inserted into plasmid pHEN6 ^58^ resulting in construct pHEN6-ADAH11 comprising a PelB secretion signal at the N-terminus and a hexa-histidine and triple-FLAG tag on the C-terminus. Nanobody was expressed in *Escherichia coli* (*E. coli*) TG1 cells cultured in 2xYT media, induced with 1mM Isopropylthio-β-galactoside (IPTG) overnight (∼16 hours) at 30°C, harvested by centrifugation and pellets stored at −80°C. Cell pellets were resuspended in ice cold TES (50 mM TRIS pH 8.0, 20% w/v sucrose, 1 mM EDTA) and incubated for 1h at 4°C. Next, Shock Buffer (20□mM Tris pH 8.0, 5□mM MgCl_2_) was added followed by incubation for 1h at 4°C. Supernatant was cleared by centrifugation, applied to 1 ml HisPur Ni-NTA resin (ThermoFisher Scientific) and incubated for 1h at 4°C with agitation. After washing with Wash Buffer (50 mM HEPES pH 8.0, 200 mM KCl, 10 mM Imidazole), ADAH11 was eluted with Elution Buffer (150 mM imidazole 50 mM HEPES pH 8.0, 200 mM KCl). Fractions containing ADAH11 protein were pooled, dialysed into PBS and further purified by SEC using a S200 10/30 GL column (Cytiva) equilibrated in PBS. Eluted ADAH11 was concentrated to 1 mg/ml, and stored at −80°C.

#### Gigabody preparation

ADDomer derived from chimpanzee adenovirus AdY25 comprising an A57S mutation (Supplementary Table 1) was expressed and purified as described above for ADDoCoV. Following SEC and IEX, samples were sterile filtered, flash frozen in liquid nitrogen and stored at −80°C. The integrity of the final sample was confirmed using both reducing SDS-PAGE and negative stain EM.

A codon optimised synthetic DNA encoding fibre tail peptide (Supplementary Table 5), T4-foldon trimerisation domain and ADAH11 spaced by glycine-serine linker sequences (Supplementary Table 6) was inserted into the pHEN6 plasmid (Genscript), expressed in T7 Express *E. coli* cells (New England Biolabs) cultured in Terrific broth (TB) medium and induced with 1mM IPTG for overnight expression at 16°C. Cells were harvested by centrifugation (4,000g for 10 min), resuspended in Lysis Buffer (50mM Tris-HCl pH 8, 300 mM NaCl, 10 mM Imidazole, 0.5mg/ml Lysozyme), frozen at −20°C and thawed at 37°C for 10 min, followed by DNase treatment at 4°C (15 min). DNase treated sample was sonicated at 50% amplitude 4 times for 30 seconds, Pulse 1s/1s using Vibracell VC 750 (Sonics and Materials) and clarified by centrifugation (12,000g for 15 min). Cleared supernatant was loaded onto a 5ml Histrap FF crude Ni-NTA affinity column (Cytiva), washed with Wash Buffer (50 mM Tris-HCl pH 8, 300 mM NaCl, 50 mM Imidazole), and nanobody trimers eluted with Elution Buffer (50 mM Tris-HCl pH 8, 300 mM NaCl, 250 mM Imidazole). Elution fractions were pooled and concentrated to 500 μl using a 10 kDa MWCO Amicon centrifugal filter unit (EMD Millipore), and further purified by SEC using a Superose 6 HR 10/30 column (Cytiva) equilibrated with PBS. Peak fractions were pooled, aliquoted and stored at 4°C.

Gigabody was assembled by mixing purified ADAH11-Trimer and AdY25 A61S ADDomer in PBS at a molar ratio of 1:1.2 pentons to ADAH11-Trimer. After 1-hour incubation rotating at 4°C, the mixture was subjected to SEC on a Superdex 200 10/300 GL column (GE Healthcare) equilibrated in PBS. Peak fractions containing Gigabody were pooled, concentrated using a 100 kDa MWCO Amicon centrifugal filter unit (EMD Millipore) and used fresh, or flash-frozen in liquid N_2_ for storage at −80°C.

### Thermostability measurements

Thermal shift experiments were performed using a ThermoFluor assay as described previously ^28^.

### Negative-stain sample preparation and electron microscopy

#### ADDoCoV

4 μl of 0.1 mg/ml ADDoCoV protein sample dialyzed into 25 mM HEPES pH 7.5, 150 mM NaCl, 2 mM EDTA was applied onto a freshly glow discharged (1 min at 10 mA) CF300-Cu grid (Electron Microscopy Sciences), incubated for 1 min, and manually blotted. 4 μL of 3% Uranyl Acetate was applied onto the same grid and incubated for 1 min before the solution was blotted off. Images were acquired at a nominal magnification of 49,000x on a FEI Tecnai 12 120 kV BioTwin Spirit microscope equipped with an an Eagle 4k x 4k CCD camera.

#### ADDoCoV-ADAH11 complex

5 μl of 0.1 mg/ml ADDoCoV-ADAH11 complex sample was prepared as above. Images were recorded at 62,000x magnification corresponding to a pixel size of 1.63 Å/pix. A total of 5,025 particles from 498 images were picked and reference free two-dimensional classification was performed leading to 1,396 particles included in final 2D class averages (Supplementary Fig. 6).

### Cryo-EM sample preparation and data collection

#### ADDoCoV

4 μl purified ADDoCoV (0.54 mg/ml) was applied to glow-discharged holey Quantifoil R 1.2/1.3 holey carbon grids (Agar Scientific), blotted for 2 seconds at 100% relative humidity and 4°C inside a Vitrobot Mark III, before plunge-freezing in 37% ethane-propane at liquid nitrogen temperature. Cryo-EM data were collected at 200 kV with a FEI Talos Arctica microscope equipped with a Gatan K2 direct electron detector and an energy filter at 20 eV slit width, using automated acquisition software (EPU). A total of 1375 dose-fractionated movies each containing 40 frames (0.2 s per frame) with an accumulated total dose of 44 e−/Å^2^ were recorded in counted super-resolution mode at a nominal magnification of 130,000x corresponding to a physical pixel size of 1.05 Å and a virtual pixel size of 0.525 Å using a defocus range of −0.7 to −2.2 μm (Supplementary Figs 2,3, Supplementary Table 2).

#### ADDoCoV-ADAH11 complex

3 μl of 1.2 mg/ml ADDoCoV-ADAH11 complex was loaded onto a glow discharged Quantifoil R1.2/1.3 holey carbon grid (Agar Scientific). The sample was incubated for 30 s at 90 % relative humidity and 16°C inside Leica EM ACE 600 (Leica EM GP2 plunge freezer), blotted for 1.2 s and vitrified in liquid ethane at liquid nitrogen temperature. Data were acquired on a FEI Talos Arctica as described above. Data were collected in counted super-resolution mode at a nominal magnification of 130,000x with a physical pixel size of 1.05 Å/pix and a virtual pixel size of 0.525 Å/pix. The total dose of 55.6 e/Å^2^. Each movie was fractionated in 45 frames of 200 ms. 11,800 micrographs were collected with a defocus range comprised between −0.8 and −2.0 µm.

### Cryo-EM data processing

#### ADDoCoV

Image processing was performed with the RELION 3.1 software package ^59^. The micrographs were motion corrected using MotionCor2 ^60^ and contrast transfer function (CTF) information determined using gctffind4.1 ^61^. 1375 micrographs with CTF rings extending beyond 4 Å were selected for further processing. 96,456 particles were boxed using RELION auto-picking software. 2D classification (Supplementary Fig. 2) and 3D classification with imposed icosahedral symmetry was performed, followed by initial 3D-autorefinement. Further rounds of 3D-classification/refinement were carried out on 32,227 polished particles after CTF refinement and spherical aberration correction before using post-processing for masking and automatic B-factor sharpening. The resolution of the final map was determined to be 2.36 Å based on the Fourier Shell Correlation (FSC) = 0.143 criterion (Supplementary Fig. 3). Local resolution was calculated using local resolution estimation programme in RELION (Supplementary Fig. 3b). 3D classification was performed using public cloud resources provided by Oracle Cloud Infrastructure as described previously ^28^.

#### ADDoCoV-ADAH11 complex

11,283 dose-fractionated movies were image processed as described above. 68,258 particles were automatically picked using Relion 4.0 ^62^. After three rounds of 2D classification, a total of 46,223 particles were selected for further 3D classification. The initial 3D model was filtered to 60 Å during 3D classification using 8 classes. The best class of 13,950 particles was selected for the following 3D-autorefinement leading to a reconstruction of ∼4.35 Å resolution. Subsequently, the maps were subjected to local defocus correction and Bayesian particle polishing in Relion 4.0. Global resolution and B-factor (−79.66 Å^2^) of the maps were estimated by applying a soft mask around the protein density using the gold-standard FSC criterion 0.143, resulting in an overall resolution of 4.06 Å (Supplementary Fig. 6). Local resolution maps were generated using Relion 4.0. The refined particles stack was expanded 60-fold according to icosahedral symmetry. The symmetry expanded particle stack was then used as input for the masked 3D classification with the focus mask corresponding to one penton base protein and the ADAH-11 region created in UCSF Chimera ^63^. The masked 3D classification was performed with 8 classes resulting three good classes with densities for ADAH11 and penton base protein (Fig. 2e.f, Supplementary Fig. 6).

### Cryo-EM model building and analysis

Homology modelling was performed using iTasser ^64^ starting from the human ADDomer structure (PDB ID 6HCR) ^28^. Using COOT ^65^, the model was fitted manually into the EM map, followed by iterative positional and B-factor refinement using Phenix Real-Space software ^66^. After adjustments in COOT the model was evaluated using Molprobity ^67^.

### *In vitro* selection of specific nanobodies by Ribosome Display

Ribosome display *in vitro* selection using a synthetic nanobody library was performed against ADDoCoV (comprising the SARS-CoV-2 S RBM AH epitope) as described ^31^. After five cycles of ribosome display against ADDoCoV immobilised on 96-well microtiter plates, the DNA pool was cloned into pHEN6. Individual colonies were picked, and nanobodies expressed in *E. coli* TG1 (Agilent Technologies) in dYT medium at 30°C overnight after induction with 1 mM IPTG. Nanobodies binding ADDoCoV, the SARS-CoV-2 RBD and S, but not ADDomer alone or a mutant S devoid of AH (SpikeΔAH), were identified by ELISA and then sequenced (Supplementary Table 3).

### Surface plasmon resonance (SPR) experiments

#### ADAH11 nanobody binding to SARS-CoV-2 RBDs

Interaction experiments using surface plasmon resonance (SPR) of ADAH11 nanobody monomer and different RBDs were carried out with a Biacore T200 system (GE Healthcare) according to the manufacturer’s protocols and recommendations. Briefly, biotinylated RBD proteins were immobilised on streptavidin-coated SA sensor chips at ∼3845 response units (RU) for Wuhan RBD and ∼2500 RU for Delta and Omicron RBD. Binders were diluted to the concentrations indicated (Fig. 2g) and passed over immobilised RBDs at a flow rate of 30 µl/minute. The Running Buffer for all SPR measurements was PBS. The sensorgrams were analyzed using the Biacore Evaluation Software (GE Healthcare) and *k*_on_, *k*_off_ and K_D_ values were determined using a two state reaction binding model. All experiments were performed in triplicates.

#### Gigabody binding to Wuhan RBD

Purified, biotinylated Wuhan RBD ligand was immobilised on a streptavidin-coated (SA) sensor chip (GE Healthcare) at 2453 RU. For all interaction measurements, the analyte was injected at a flow rate of 50 μl/min for 120s using PBS as the Running Buffer. Dissociation was performed for 600s. Gigabody, and AdY25 ADDomer as a negative control, were serially diluted and injected at concentrations of 0.5nM, 1.0nM, 1.5nM and 2.0nM. The chip was regenerated using 2 injections of 10mM glycine pH 2.6. All measurements were performed in triplicates. Final sensorgrams were obtained by subtracting the control sensorgrams from the corresponding Gigabody sensorgrams accounting for non-specific binding to the sensor chip. Fitting with Biacore Evaluation Software (GE Healthcare) indicated picomolar binding (K_D_=30±20pM) dictated by very slow dissociation kinetics.

#### Gigabody binding to Omicron RBD

Purified, biotinylated Omicron RBD ligand was immobilised on a SA sensor chip (GE Healthcare) at 3622 RUs. Injection, dissociation and regeneration were performed as above, for Gigabody serial diluted at concentrations of 1.0nM, 1.5nM, 2.0nM and 2.5nM. Sensorgrams were analyzed with the Biacore Evaluation Software, again indicating picomolar binding (K_D_=10±3pM) with very slow dissociation kinetics, similar to Wuhan RBD.

### Molecular Dynamics simulations

#### Construction of a complete ADDoCoV model using Rosetta and MD

The input model was based on the cryo-EM structure combined with the AH-epitope sequence (Fig. 1d, Supplementary Table S1) added manually adopting a structure derived from the ACE2 receptor in complex with the RBD of the S protein (PDB ID 7C8D) ^68^. The RGD loop, unresolved in the cryo-EM density, was reconstructed using Rosetta ^69–73^. Symmetrical pentamer models were generated with Rosetta SymDock ^71, 72^, and the introduced 5-fold symmetry was maintained during all following steps. Missing loops were reconstructed using Rosetta Remodel ^69, 73^. Models were relaxed using Rosetta Relax ^70^, and subjected to MD simulations with GROMACS 2019 ^74^. The ADDoCoV structure was parametrised with the gromos54a7 forcefield in a cubic box with simple point charge water and sodium ions to neutralize the net charge. MD comprised 5 replicates of 100 ps of NVT followed by 67 ns of NPT simulations. Trajectories were analyzed with CPPTRAJ ^75^. All analyses were based on Cα positions if not stated otherwise. The first 10□ns of each production MD run were excluded from all analyses to allow time for system equilibration. The conformational landscape of the AH epitope was analyzed by principal component analysis (mdtraj ^76^ and sklearn ^77^) and cluster analyses (cpptraj, kmeans algorithm) based on the same cartesian space.

#### Construction of a Gigabody model using Rosetta and MD

Missing loops in the ADDomer model, sequence adjusted for the AdY25 penton base protomer (Supplementary Table 1), were constructed using Rosetta SymDock ^71, 72^, Remodel ^69, 73^ and Relax ^70^ as described for the ADDoCoV model. The trimeric ADAH11 nanobody fibre tag structure was modeled based on the bacteriophage T4 fibritin derived trimeric foldon structure (PDBID: 4NCV) ^78^. The structures of the fibre tail peptide fused to the N-terminus of the foldon, and ADAH11 nanobody fused to the C-terminus, were predicted with trRosetta ^79^. Two amino acids of the foldon were included during the prediction with trRosetta at the C-terminus of the fibre tail peptide and the N-terminus of the nanobody, respectively (Supplementary Fig. 8). The fibre tail peptides and nanobodies were aligned to the trimeric foldon using these overlapping residues. The complete ADAH11-Trimer structure was subsequently relaxed with Amber ^75^ using the ff14SB forcefield and no solvent in three consecutive minimisations with 1000 cycles with positional restraints on all atoms of 10, 1, and 0 kcal/mol/A^2^.

Next, the structure was further relaxed with Rosetta ^70^ using the MonomerRelax2019 script and the 10 lowest energy structures were placed manually in PyMol on top of the AdY25 ADDomer A57S model guided by the Ad5 penton base fibre tail peptide complex (PDB ID 1X9T) ^35^. Subsequently, Amber ^75^ was used to relax the fibre tail peptide to the position observed in the experimental structure (PDB ID 1X9T) ^35^. 50 ns MD simulations were then performed with Amber. The protein was parametrised with the united-atom forcefield ff03u ^80^. The ADDomer-fibre tail peptide complex was relaxed with increasing positional restraints (1000 steps, ntmin = 3, restraint_wt = 0.1, 0.2, 0.5, 1, 2, 5, and 10) on the ADDomer and the fibre tail peptides to generate a complex in the experimentally-observed conformation (PDB ID 1X9T) ^35^.

Two additional minimisation steps were performed, the first without positional restraints and the second with implicit Born solvation model (IGB=1) ^81^. After an initial heating step (0.05 ns from 0.1 to 300 K), 50 ns MD simulations were performed with 2 fps timestep using implicit solvation and Langevin dynamics (ntt=3) for each of the 20 starting structures (10 Rosetta models * 2 conformations). Four runs with the three fibre tail peptides in proximity and three runs with two peptides in proximity were unstable and discarded. To illustrate the full scale of the Gigabody (Fig. 3c, Supplementary Fig. 8), the modeled pentamers from these simulations were aligned with the pentamers in the ADDomer cryo-EM structure (PDB ID 6HCR) ^28^.

### SARS-CoV-2 MiniV preparation

Artificial minimal SARS-CoV-2 virions (MiniVs) were assembled from small unilamellar vesicles (SUVs) as described previously ^36^. Briefly, SUVs containing NTA(Ni^2+^) and Rhodamine B-functionalised membranes were coupled to recombinant Wuhan SARS-CoV2 S ectodomains bearing an oligohistidine tag ^34^. SUVs were prepared by membrane extrusion to obtain a monodisperse vesicle population with a mean diameter of 100 nm from a lipid solution of 45□mol% DOPC, 21□mol% DOPE, 3□mol% DOPS, 12□mol% DOPI, 14□mol% cholesterol, 3□mol% SM, 1 mol% DGS-NTA(Ni^2+^) and 1 mol% Rhodamine B-PE (all lipids obtained from Avanti Polar Lipids).

MiniV size distribution was measured by dynamic light scattering with a Malvern Zetasizer Nano ZS system at a total lipid concentration of 100□µM in PBS. Temperature equilibration time was set to 300□s at 25□°C, followed by three repeated measurements for each sample at a scattering angle of 173° using the built-in automatic run-number selection. The material refractive index was set to 1.4233 and solvent properties to η□=□0.8882, n□=□1.33 and ε□=□79.0. For assessment of Gigabody-mediated MiniV clustering, the MiniV solution was preincubated with 1.5 nM Gigabody for 30 min in the dark at 4°C before measurement.

For confocal microscopy observation of MiniV-cell attachment after 2.5 hours of incubation under control conditions, or with addition of 500 nM nanobodies or 1.6 nM Gigabody, respectively, A549 cells stably expressing ACE2 ^82^ were stained with CellTracker Green CMFDA dye (Invitrogen, USA) according to the manufacturer’s recommendations. Nuclei were stained with 10 µM Hoechst33342 (Sigma Aldrich). Laser scanning confocal microscopy was performed with a LSM 800 (Carl Zeiss AG). Images were acquired with a ×63 immersion oil objective (Plan□Apochromat ×63/1.40 Oil DIC, Carl Zeiss AG). Analysis was performed with ImageJ (NIH) and adjustments to image brightness and contrast, as well as background corrections, were always performed on the whole image and special care was taken not to obscure or eliminate any information from the original image.

### SARS-CoV-2 MiniV retention assays

Retention assays were performed as described previously ^36^ using human ACE2 expressing A549 cells. Briefly, MiniVs were incubated with A549 cells at a final lipid concentration of 10 µM in flat bottom 96 well plates and in low serum containing culture medium (DMEM supplemented without phenol red, 4.5□g/l glucose, 1% L-glutamine, 1% penicillin/streptomycin, 0.01□mg/ml recombinant human insulin, and 0.5% fetal bovine serum). After 2.5 hours, MiniV Rhodamine B fluorescence was measured with a plate reader for each well in 4 positions. Cultures were afterwards washed 3 times with PBS. Subsequently, residual fluorescence was measured in each well and normalised to the initial fluorescence intensity to calculate MiniV retention values after correction for background fluorescence and negative controls. Gigabody dilution curves for retention analysis were prepared by preincubating MiniVs with 1.6 nM Gigabody for 30 min at 4°C in the dark before addition to the cells.

### Mouse immunisation experiments

Female C57BL/6 mice were obtained from Charles River Laboratories (UK) and maintained at the University of Bristol Animal Services Unit in specific pathogen-free conditions in accordance with established practices and under a UK Home Office License ^83^. Mice were immunised with 40μg ADDoCoV vaccine or ADDomer scaffold as a control via intranasal, intramuscular or subcutaneous routes (n = 10 mice per treatment group pooled across 2 experimental replicates) on day 0 (primary immunisation), day 21 (boost 1) and day 42 (boost 2). Mice were humanely euthanised on day 62; 9 weeks post initial immunisation, by terminal exsanguination under general anesthesia.

#### Intranasal (IN)

Mice were lightly anaesthetised using isoflurane, and 12.5 μL ADDoCoV vaccine in sterile PBS (1.6 mg/ml) was instilled into each nostril (total dose 25 μL; 40 μg).

#### Intramuscular (IM)

Mice were lightly anaesthetised using isoflurane and received intramuscular injection with 50μL ADDoCoV vaccine in sterile PBS (0.8 mg/ml) into the quadriceps muscle using a 25G 5/8 inch needle (total dose 50 μL; 40 μg).

#### Subcutaneous (SC)

Non-anaesthetised mice were restrained in a tube restrainer. Subcutaneous injection was performed with 50μL ADDoCoV vaccine in sterile PBS (0.8mg/ml) using a 25G 5/8 inch needle at the tail base (total dose 50 μL; 40 μg).

#### Sample collection

The presence of serum antibody was assayed in peripheral blood at baseline (day-1), day 20, and day 41 and in terminal bleeds on day 62. Peripheral blood samples (30-50μL) were collected from the lateral tail vein. For collection of terminal blood samples, mice were deeply anesthetised using isoflurane, and 500-800μL of blood was collected following thoracotomy and cardiac puncture. Peripheral blood and terminal bleed samples were processed for serum collection. Blood was collected into autoclaved microcentrifuge tubes without anti-coagulant and allowed to clot at room temperature for 20 min. Samples were centrifuged at 2000 g for 10 min at 4°C. Serum was transferred to a fresh microcentrifuge tube, and centrifugation was repeated at 2000 g for 10 min at 4°C. Following centrifugation, serum was transferred to a fresh microcentrifuge tube and frozen in aliquots at −80°C. Nasal washes (NW) and bronchoalveolar lavages (BAL) were taken post-mortem using established methodology ^84, 85^.

The presence of mucosal antibody in murine nasal secretions were assayed by flushing a 500μL volume of ice-cold PBS through the nasal turbinates. Briefly, scissors were used to make an incision from the abdomen to the jaw in order to expose the thoracic cage and neck. The trachea was exposed and a 20G x32mm Surflo intravenous catheter (VWR international) inserted. A 1 ml syringe containing 500μL PBS was then attached and the fluid used to flush the nasal cavity. Fluid existing the nares was captured using an Eppendorf and then incubated on ice with Protease inhibitor cocktail (Roche Diagnostics). Washes were centrifuged at 1000 g for 10 min at 4°C to remove cellular debris and mucus ^86^. Fluid supernatants were transferred to fresh autoclaved microcentrifuge tubes and immediately frozen in aliquots at −80°C.

To isolate mucosal antibody in the lower respiratory tract, lung lavages were performed. Briefly, a 20G intravenous catheter with stylet withdrawn was inserted, directed towards the lungs. A syringe containing 1 ml ice-cold PBS was used to aspirate the lungs, ensuring not to overinflate and rupture the tissue. To prevent PBS from leaking from the catheter insertion site, thread was used to tie off the catheter to the trachea. Recovered PBS from lung washes were again incubated on ice with protease inhibitor then centrifuged at 1000 g for 10 min at 4°C to remove cellular debris and mucus^85^. Fluid supernatants were transferred to fresh autoclaved microcentrifuge tubes and immediately frozen in aliquots at −80°C.

### Enzyme–linked immunosorbent assay (ELISA)

#### Antigen-specific serum antibody ELISA

The antigen-specific IgG and IgA titres in mouse sera were assessed by a semi-quantitative ELISA. MaxiSorp high binding ELISA plates (Nunc) were coated with 100□μl/well of 1□μg/ml highly purified SARS-CoV-2 SARS-CoV-2 RBDs. For the IgG and IgA standards, plates were coated with 1:1000 dilution each of goat anti-mouse Kappa (Catalog #1050-01, Southern Biotech) and Lambda light chains (Catalog #1060-01, Southern Biotech). After overnight incubation at 4□°C, the plates were washed 4 times with PBS-Tween 20 0.05% (v/v) and blocked for 1□h at 37□°C with 200□μl/well blocking buffer (1% BSA (w/v) in PBS-Tween-20 0.05%(v/v)). The plates were then washed, and 10-fold serial dilutions of serum samples (10^3^-10^6^), or a 5-fold dilution series starting at 200ng/ml of purified IgG (Catalog #0107-01, Southern Biotech) or IgA (Catalog #0106-01, Southern Biotech) were added using 50□μl/well volume. Plates were incubated for 1□h at 37□°C, then washed and secondary antibody added at 1:2000 or 1:4000 dilution in blocking buffer (100□μl/well) using either anti-mouse IgG-HRP (Catalog #1030-05, Southern Biotech), or anti-mouse IgA-biotin (Catalog #1040-80, Southern Biotech). After a 1h incubation at 37°C, plates incubated with biotinylated antibody were washed and incubated at 37°C for 1 hr with a 1/200 dilution of Streptavidin-HRP (Catalog #890803, R&D systems). Plates were then washed and developed using 100□μl/well SureBlue TMB (3,3′, 5,5′-tetramethylbenzidine) substrate, and the reaction stopped after 5□min with 100□μl/well stop solution (Insight Biotechnologies). The absorbance was read on a FLUOstar Omega multi-mode microplate reader at 450□nm (BMG LABTECH). For assaying mucosal samples for the presence of antigen-specific IgG and IgA antibody, the same procedure was followed except mucosal samples were used at a 1/10 - 1/250 dilution series. To determine the presence of cross-reactive antibody in the serum or mucosal secretions of vaccinated mice, the binding of antibody to different variants of SARS-CoV-2 RBD (Wuhan, Alpha, Beta, Delta and Omicron) was measured. Here, the variant RBDs were used to coat MaxiSorp high binding plates and ELISA performed as before.

#### Nanobody ELISA

Proteins ADDomer, ADDoCoV, RBD, S, SpikeΔAH and BSA were produced and purified as described ^57^. Highly purified proteins were diluted in PBS to a final concentration of 40 µg/ml. Next, 100 µl of the diluted proteins were added to the corresponding well in a microtiter plate followed by a gentle tap to ensure even coating of all wells before sealing the plate and incubating overnight at 4°C. On the following day, supernatants were discarded and the plate washed 3 times with 300 µl Wash Buffer (PBS pH 7.4, 0.1% Tween) before drying the plate by placing it upside down on a paper towel to remove residual Wash Buffer. Next, 200 µl Blocking Solution (PBS pH 7.4, 5% milk) was added to each well, and the plate was incubated at room temperature for 1 hour. The nanobody samples were diluted in Wash Buffer to a concentration of 1μM concentration. Subsequently, the Blocking Buffer was removed from plate before drying on a paper towel, followed by adding 100 µl of corresponding nanobodies or control buffer, respectively, to each well of the plate before incubating the plates at room temperature for 1 hour. Next, the samples were removed from the plate, and the plate was washed with 300 µl Wash Buffer 3 times and then dried on paper towel. Finally, 50 µl of anti FLAG-HRP antibody (dilution 1:3000) was added to each well, and the plate was incubated at room temperature for 1 hour. Sample was removed, and the plate was washed with 300 µl Wash Buffer 3 times and dried on a paper towel. Then, 100 µl of TMB reagent was added to each well, followed by incubation at room temperature for 5 min and stopping the reaction with addition of 50 µl of 1N HCl to each well. Finally, absorbance at 450 nm was measured in a microplate reader. The data was plotted using Microsoft excel. The standard deviation of triplicates was added as error bars.

### SARS-CoV-2 virus neutralisation assay

Vero E6 cells engineered to express the cell surface protease TMPRSS2 (VeroE6-TMPRSS2) ^87^ (NIBSC) and Caco-2 cells engineered to express ACE2 ^88^ were cultured at 37°C in 5% CO_2_ in DMEM containing GlutaMAX (Gibco, Thermo Fisher) supplemented with 10% (v/v) FBS (Gibco) and 0.1 mM non-essential amino acids (NEAA, Sigma Aldrich). The ADAH11 nanobody was serially diluted 2-fold for eight dilutions, from a 0.85 µg/ml starting dilution, in triplicate, in Minimum Essential Media (MEM, Gibco) containing 2% (v/v) FBS and NEAA. The ancestral SARS-CoV-2 isolate hCoV-19/England/02/2020 (GISAID ID: EPI_ISL_407073) was grown on VeroE6-TMPRSS2 cells and titrated as previously described ^87^. Virus (60 µl of 8 x 104 TCID50/ml) was mixed 1:1 with dilutions of ADAH11 and incubated for 60 min at 37°C. Following the incubation, supernatants were removed from Caco-2-ACE2 and VeroE6-TMPRSS2 cells seeded previously in µClear 96 well microplates (Greiner Bio-One) and replaced with 100 µl of the virus:sera dilutions followed by incubation for 18 hours at 37°C in 5% CO_2_. Control wells containing virus only (no ADAH11) as well as a positive control (a commercial monoclonal antibody (Absolute Antibody; Sb#15) recognising the S protein RBD) and media only negative control were also included on each plate. Cells were fixed by incubation in 4% paraformaldehyde for 60 min followed by permeabilisation with Triton-X100 and blocking with bovine serum albumin. Cells were stained with DAPI (Sigma Aldridge) and an antibody against the SARS-CoV-2 nucleocapsid protein (1:2000 dilution, 200-401-A50, Rockland) in combination with a corresponding fluorophore conjugated secondary antibody (Goat anti-Rabbit, AlexaFluor 568, Thermo Fisher). Images were acquired on the ImageXpress Pico Automated Cell Imaging System (Molecular Devices) using a 10X objective. Stitched images of 9 fields covering the central 50% of the well were analysed for infected cells using Cell ReporterXpress software (Molecular Devices). Cell numbers were determined by automated counting of DAPI stained nuclei, infected cells were determined as those cells in which positive nucleocapsid staining, associated with a nucleus, was detected. The percentage of infected cells relative to control wells containing virus only (no ADAH11) were calculated.

### Surrogate virus neutralisation assay (sVNT)

Remaining samples of sera from corresponding administration routes along with prebleed samples were pooled and passed through a protein A column and the recovered IgGs used in SARS-CoV-2 surrogate virus neutralisation assays (sVNT) ^38^ using a commercial kit (GenScript).

#### Statistics

Statistical significance was determined by calculating standard deviations following standard mathematical formulae. For biochemical experiments, standard deviations were calculated from independent triplicates unless indicated otherwise. For mouse immunisation data, statistical analyses were carried out using a Mann-Whitney nonparametric *t* test and GraphPad Prism software.

## Acknowledgements

We thank all members, present and past, of the Berger and Schaffitzel laboratories, and the colleagues from Bristol UNCOVER group, for their contributions and helpful discussions. We acknowledge support and assistance by the Wolfson Bioimaging Facility and the GW4 Facility for High-Resolution Electron Cryo-Microscopy funded by the Wellcome Trust (202904/Z/16/Z and 206181/Z/17/Z) and BBSRC (BB/R000484/1). We are grateful for support from the Oracle Higher Education and Research program to enable cryo-EM data processing using Oracle’s high-performance public cloud infrastructure (https://cloud.oracle.com/en_US/cloud-infrastructure), and we thank Simon Burbidge, Christopher Woods, Matt Williams and Richard Pitts for computation infrastructure support. This work was carried out using the computational and data storage facilities of the <u>Advanced Computing Research Centre</u>, University of Bristol. The authors thank University of Bristol and the Max Planck Gesellschaft (MPG), Germany, for generous support through the Max Planck Bristol Centre for Minimal Biology (MPBC).

## Author contributions

F.G., C.S. and I.B. conceived the study. F.G., D.B., K.G., J.C., C.F., A.H., G.B., S.H., M.V.V., F.D. and R.V. produced and purified protein samples, and carried out biochemical and biophysical experiments. S.K.N.Y. and U.B. prepared grids and collected EM data, S.K.N.Y. carried out image processing, D.B., A.B. and J.C.B. carried out model building and structural analysis. H.A.B., A.J.M. and J.L.R.A. performed and interpreted simulations. M.K.W. and A.D.D. performed live virus CL3 work and analyzed data. O.S. and J.S. performed and analyzed synthetic SARS-CoV-2 virion experiments. E.M., G.E., B.V.V.M., O.F., J.R., L.W., J.H., D.M., J.M., A.F. and M.B. planned, performed and analyzed mouse immunisation experiments. D.B., F.G., C.S. and I.B. wrote the manuscript with input from all authors.

## Data availability

All data needed to evaluate the conclusions in the paper are present in the paper and/or the Supplementary Materials. All datasets generated during the current study have been deposited in the Electron Microscopy Data Bank (EMDB) under accession numbers EMD-16512 (ADDoCoV), EMD-16522 (ADDoCoV-ADAH11), and in the Protein Data Bank (PDB) under accession number PBD ID 8C9N (ADDoCoV). Reagents are available from F.G., C.S. and I.B.

