## Supplementary Information for "Antibodies generated *in vitro* and *in vivo* elucidate design of a thermostable ADDomer COVID-19 nasal nanoparticle vaccine"

**Brief description of what this file includes:**

Supplementary Fig. 1: Purification and quality control of ADDoCoV.

Supplementary Fig. 2: Electron microscopy of ADDoCoV.

Supplementary Fig. 3: Quality of the ADDoCoV map and model.

Supplementary Fig. 4: ADDoCoV Dynamics.

Supplementary Fig. 5: ADAH11 nanobody binding to ADDoCoV.

Supplementary Fig. 6: Electron microscopy of ADDoCoV-ADAH11 nanobody complex.

Supplementary Fig. 7: ADAH14 and ADAH15 binding to SARS-CoV-2 Wuhan RBD.

Supplementary Fig. 8: Gigabody purification and quality control.

Supplementary Fig. 9: Gigabody modeling.

Supplementary Table 1: Cryo-EM data collection and refinement statistics, ADDoCoV.

Supplementary Table 2: Sequences of nanobodies selected by Ribosome display.

Supplementary Table 3: Cryo-EM data collection statistics, ADDoCoV-ADAH11 complex.

Supplementary Table 4: Adenoviral fiber tail peptide sequences.

Supplementary Table 5: Sequences of penton base protomers used in this study.

Supplementary Table 6: Sequences of proteins used for Gigabody preparation.

Caption for Supplementary Movie 1: Architecture of ADDoCoV nanoparticle vaccine.

Caption for Supplementary Movie 2: Gigabody nanoparticle displaying ADAH11 nanobody trimers

**Other Supplementary Materials included for this manuscript:**

Supplementary Movies 1, 2

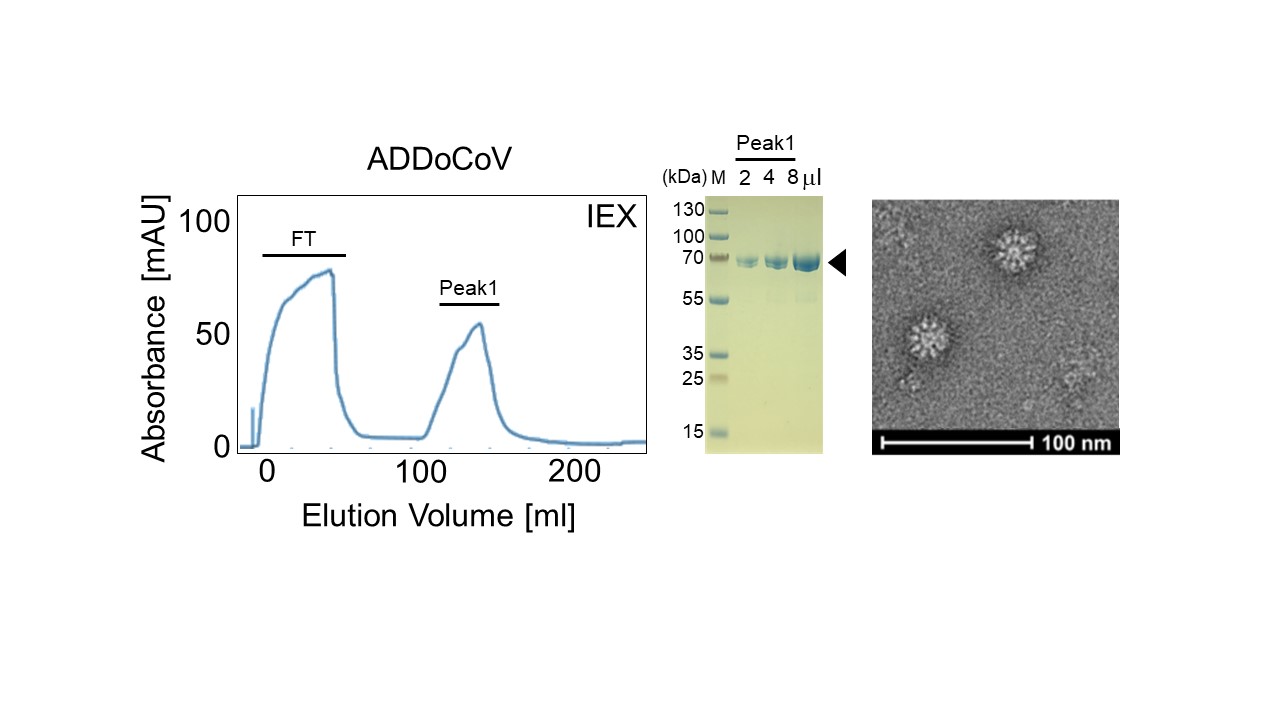

**Supplementary Fig. 1: Purification and quality control of ADDoCoV.** Ion exchange (IEX) chromatogram of ADDoCoV S protein using a Bio-Scale Mini Macro Prep High Q column (left). Absorption was detected at 280 nm (blue line). Flow through (FT) and peak fractions are indicated. ADDoCoV protein (Peak 1) was confirmed by the SDS-PAGE analysis (middle) of pooled Peak 1 fractions. Aliquot volumes loaded are indicated. M stands for molecular weight marker, molecular weights (in kDa) corresponding to marker bands are indicated. Negative-stain EM micrograph of Peak 1 is shown (right, scale bar: 100 nm). This material was used for negative-stain EM and cryo-EM sample preparations.

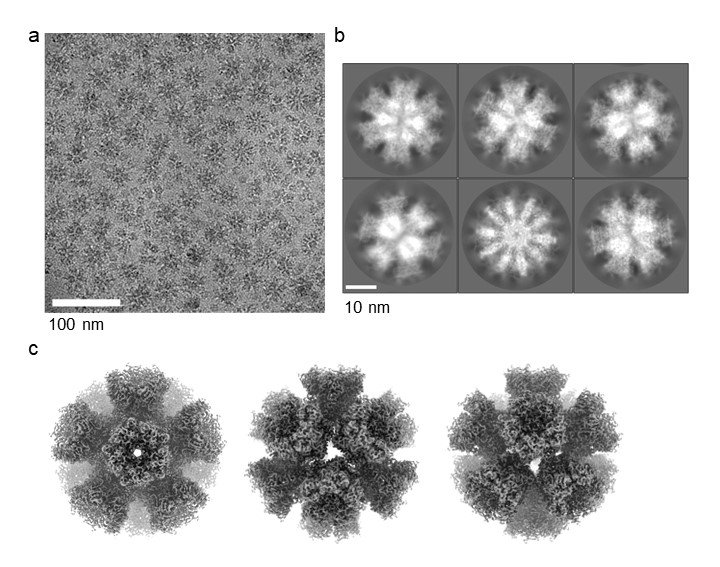

**Supplementary Fig. 2: Electron microscopy of ADDoCoV**. **(a)** A representative cryo-EM micrograph with ADDoCoV particles is shown. Some dissociation of ADDoCoV particles into pentons is observed during cryo-grid preparation. **(c)** Reference-free 2D class averages from RELION 3.1. Scale bars are indicated. **(d)** Final Cryo-EM map with icosahedral symmetry at 2.36 Å. Three different views are shown.

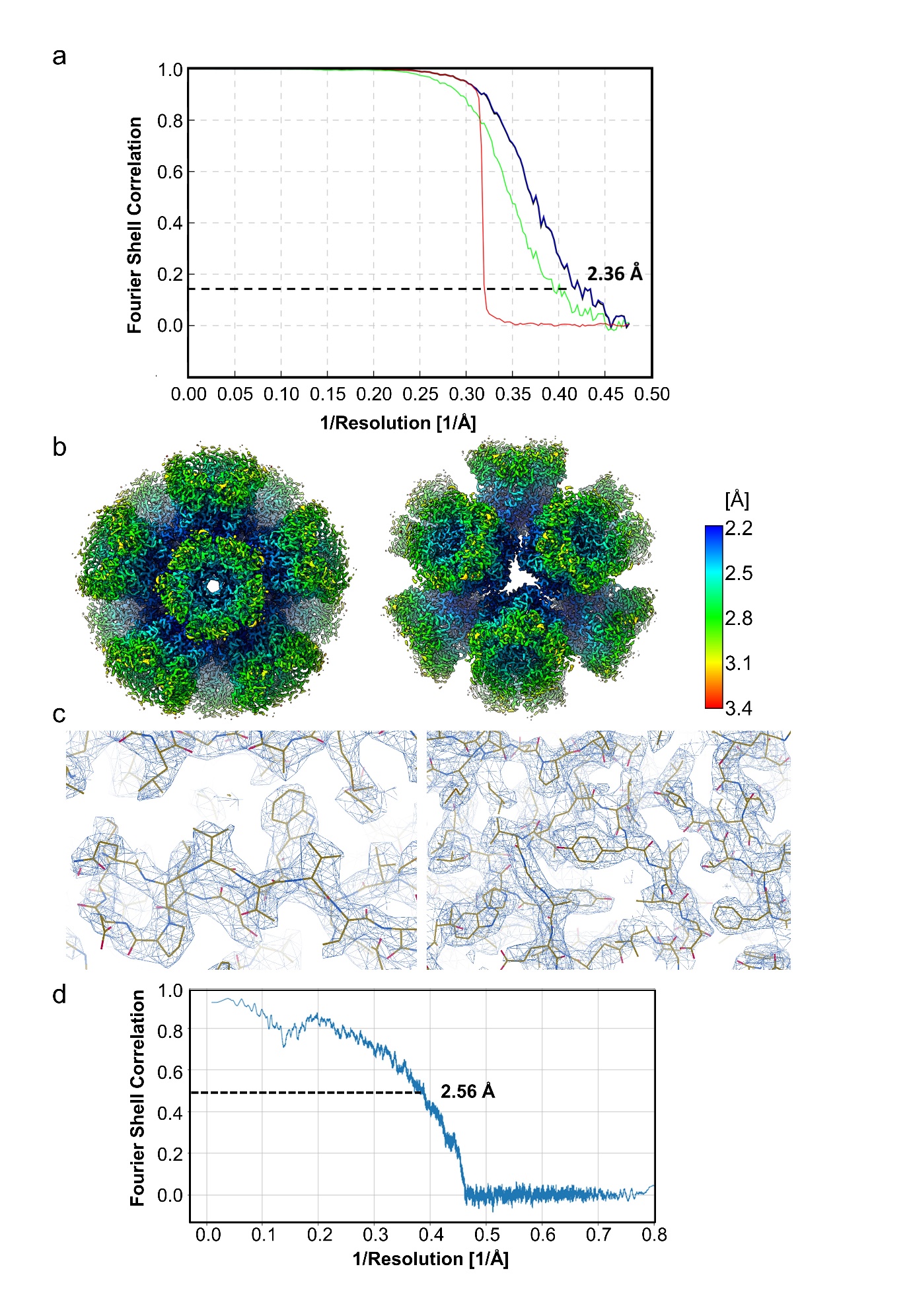

**Supplementary Fig. 3: Quality of the ADDoCoV map and model. (a)** The Fourier Shell correlation (FSC) curve after gold-standard refinement of 32,227 particles (blue curve). The FSC = 0.143 criterion indicates an overall resolution of 2.36 Å. Green curve: FSC curve of unmasked maps; red curve: FSC curve of phase randomized masked maps. **(b)** Local resolution of the final ADDoCoV cryo-EM map calculated in RELION 3.1. The core of the complex is resolved at 2.2 Å whereas peripheral parts comprising the VL and RGD loops have a lower resolution of ~ 3-3.2 Å. **(c)** Representative EM density of the ADDoCoV containing the refined atomic model. **(d)** FSC curve calculated between the atomic model and the final cryo-EM map. The map/model FSC at 0.5 reaches a resolution of 2.56 Å.

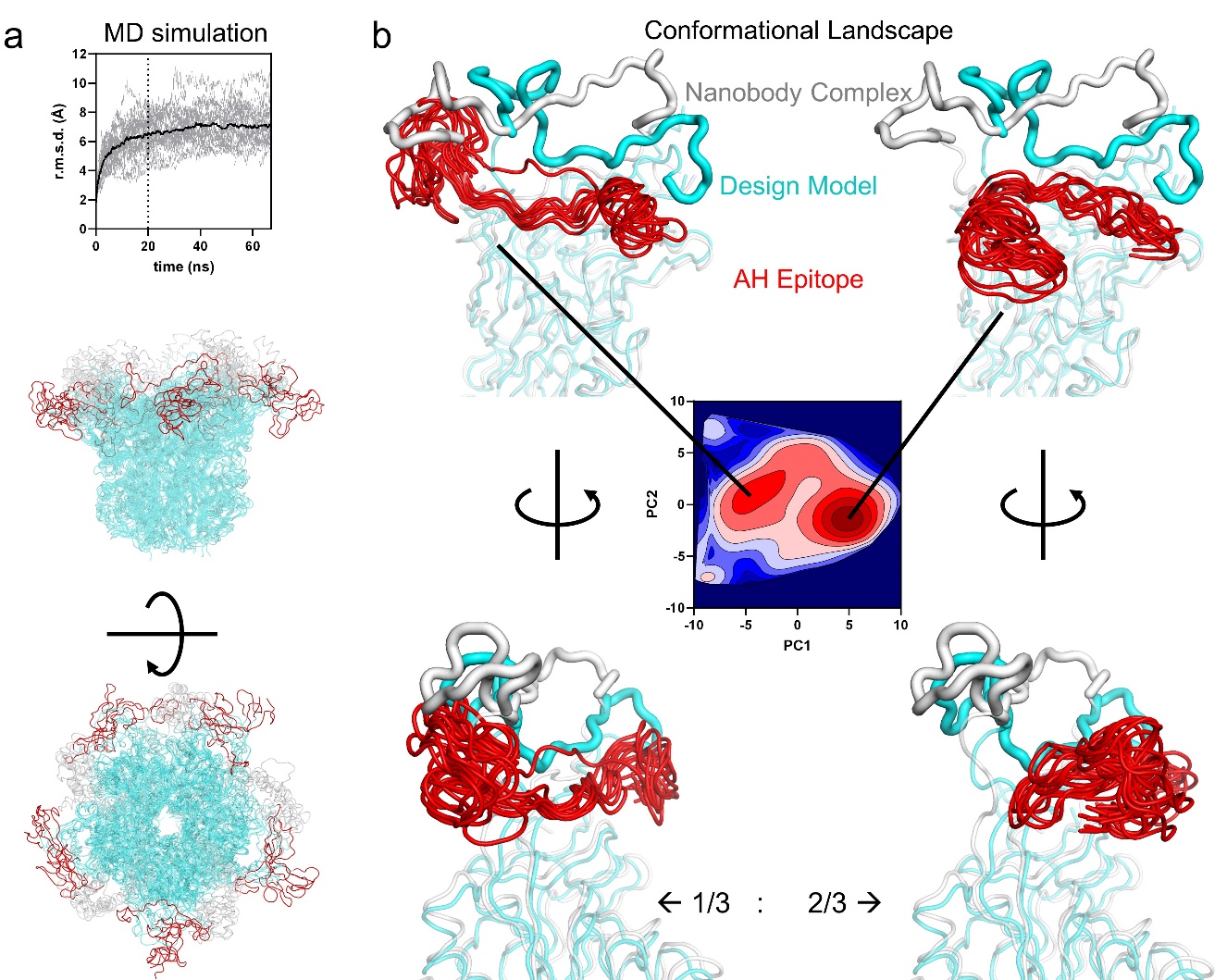

**Supplementary Fig. 4: ADDoCoV Dynamics** (**a**) MD simulations of the ADDoCoV protomer were performed based on the Rosetta Model (AH-epitope: red; RGD-loop: grey; Scaffold: cyan). As expected, the AH-epitope and RGD-loop were flexible, whereas the overall protein (cyan) remained similarly structured to its cryo-EM model. (**b**) The dynamics of the AH-epitope were analyzed for each monomer separately, resulting in an accumulated simulation time of 5 monomers x 5 replicates x 65 ns, which allowed for analysis of the conformational dynamics of the epitope in detail by principle component analysis and cluster analysis. The AH-epitope explores a broad and shallow conformational landscape with two pronounced minima. 1/3 of the trajectory populated a minimum corresponding to a conformation similar to the designed model (cyan) resembling the conformation in the open SARS-CoV-2 S form. The other 2/3 of the population corresponds to a flipped-down conformation, in which a part of the epitope protrudes sideways into the space in between pentons.

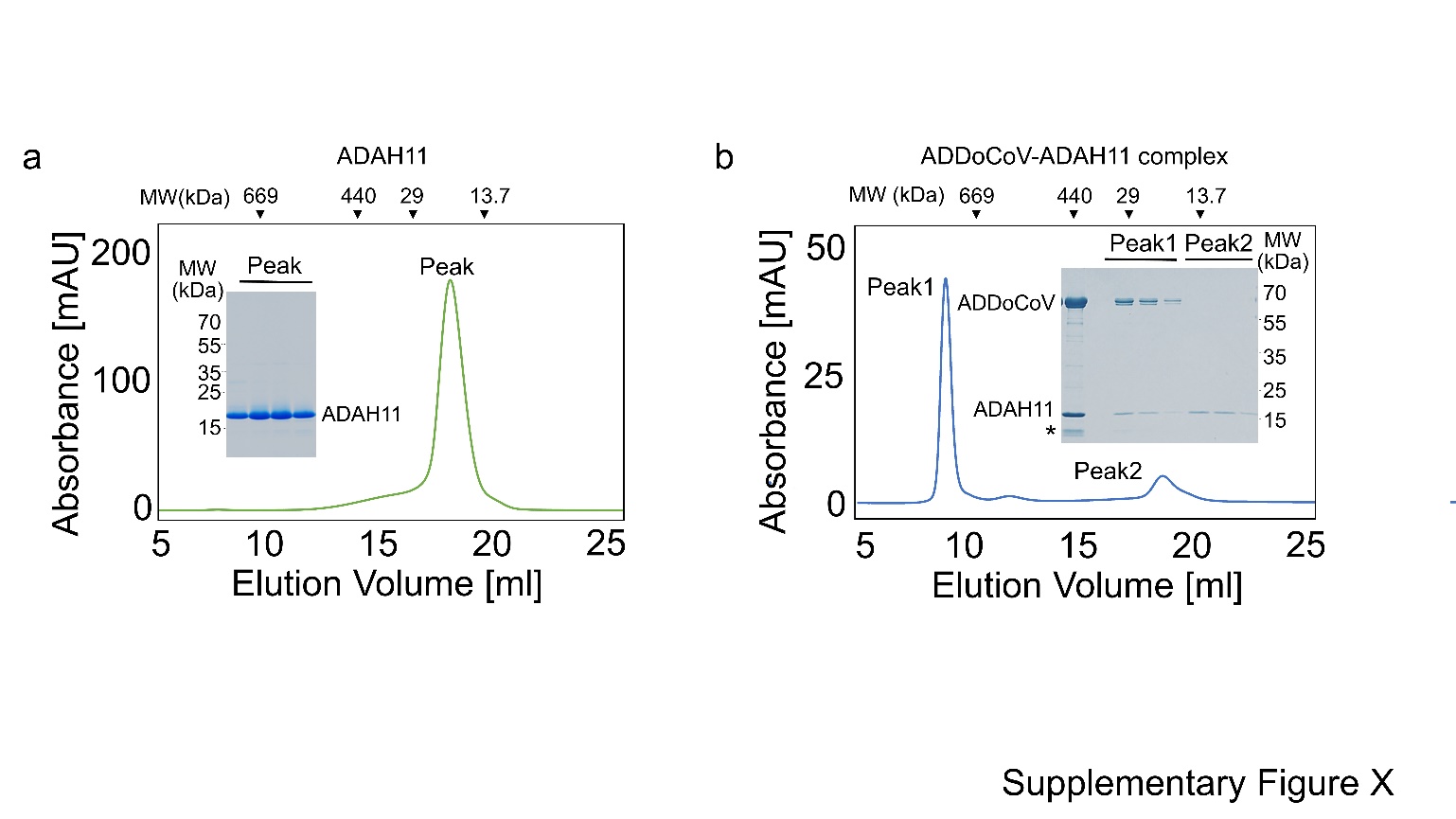

**Supplementary Fig. 5: ADAH11 nanobody binding to ADDoCoV. (a)** Size exclusion chromatography of purified ADAH11 using a Superdex 200 column is shown. Inset: Coomassie stained SDS PAGE section showing the peak fractions. **(b)** Size exclusion chromatography of ADAH11 and ADDoCoV using a Superdex 200 column. Inset: SDS-PAGE gel showing the input mixture (lane 1), Peak 1 with bands corresponding to the complex of ADAH11 and ADDoCoV and Peak 2 corresponding to excess ADAH11. The star corresponds to a degradation product of the nanobody in the input mix.

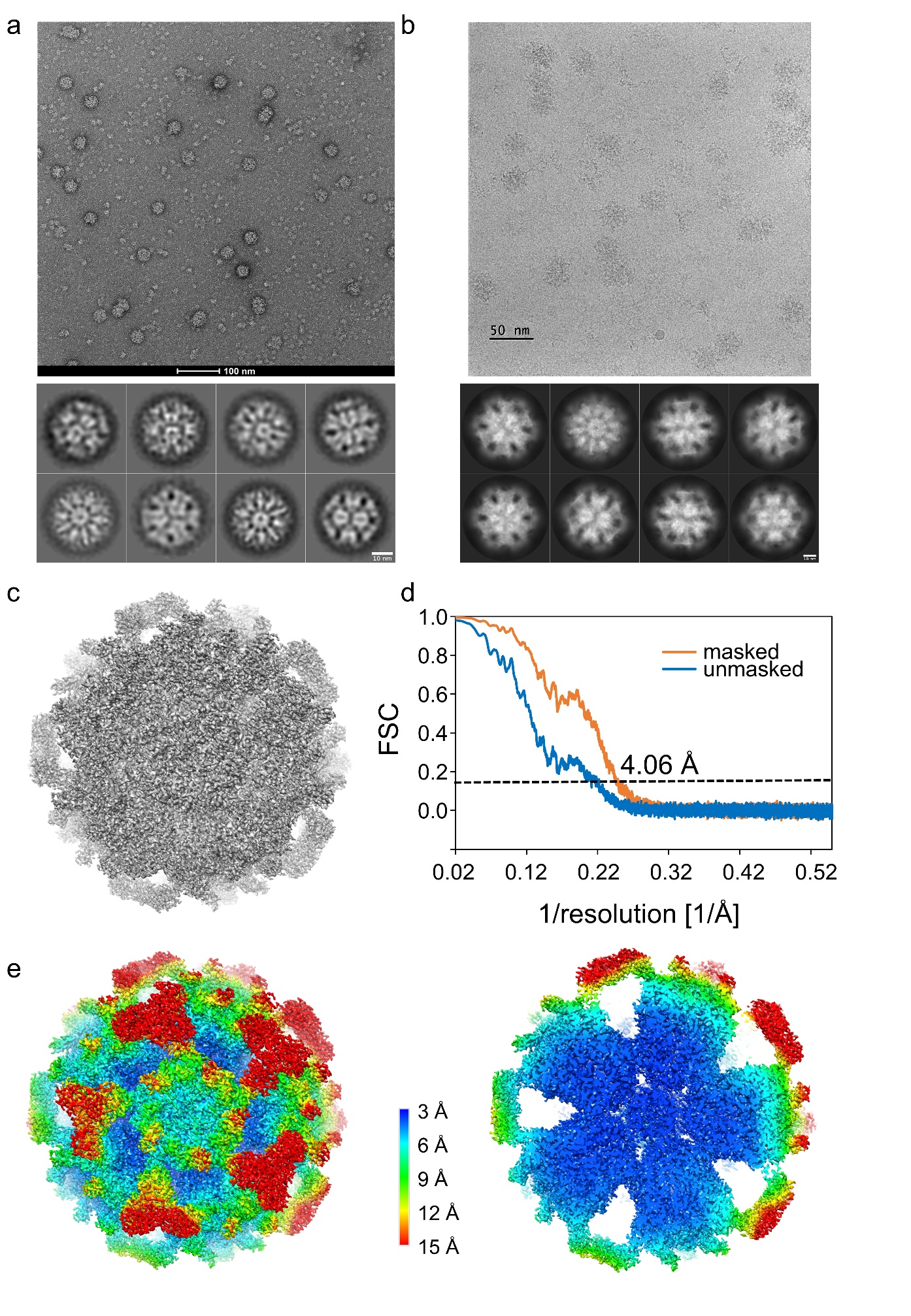

**Supplementary Fig. 6: Electron microscopy of ADDoCoV-ADAH11 nanobody complex. (a)** Negative stain micrograph showing purified ADDoCoV-ADAH11 complex and reference-free 2D class averages. Scale bars correspond to 100 nm and 10 nm, respectively. **(b)** A representative cryo-EM micrograph with ADDoCoV-ADAH11 nanobody complex and reference-free 2D class averages. Scale bars correspond to 50 nm and 10 nm, respectively. **(c)** Final Cryo-EM map of the ADDoCoV-ADAH11 complex. No symmetry was applied. **(d)** The Fourier Shell correlation (FSC) curve after gold-standard refinement. The FSC = 0.143 criterion indicates an overall resolution of 4.06 Å. Blue curve: FSC curve of unmasked maps; orange curve: FSC curve of phase randomized masked maps. **(e)** Local resolution of the final ADDoCoV-ADAH11 complex cryo-EM map calculated in RELION 4.0. The core of the complex is resolved at ~3 Å whereas peripheral parts comprising the VL loop and the bound nanobody have a lower resolution of ~ 10-15 Å which is insufficient to build an atomic model.

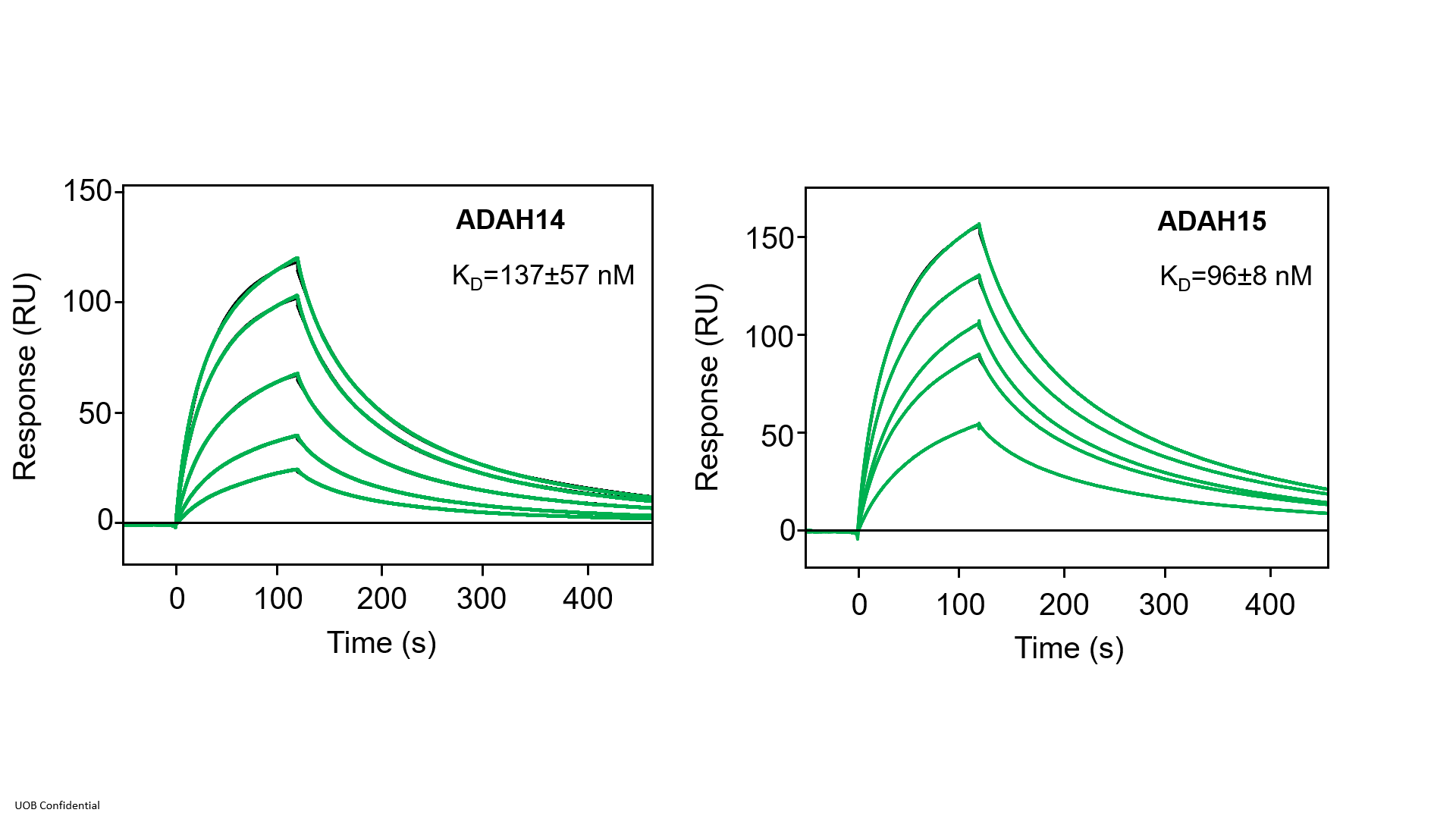

**Supplementary Fig. 7: ADAH14 and ADAH15 binding to SARS-CoV-2 Wuhan RBD.** Surface plasmon resonance of nanobodies ADAH14 (left) and ADAH15 (right) binding to immobilised Wuhan RBD. Nanobodies were injected in a concentration range from 50 nM to 250 nM. Data (green lines) were fitted (black lines) using the Biacore evaluation software.

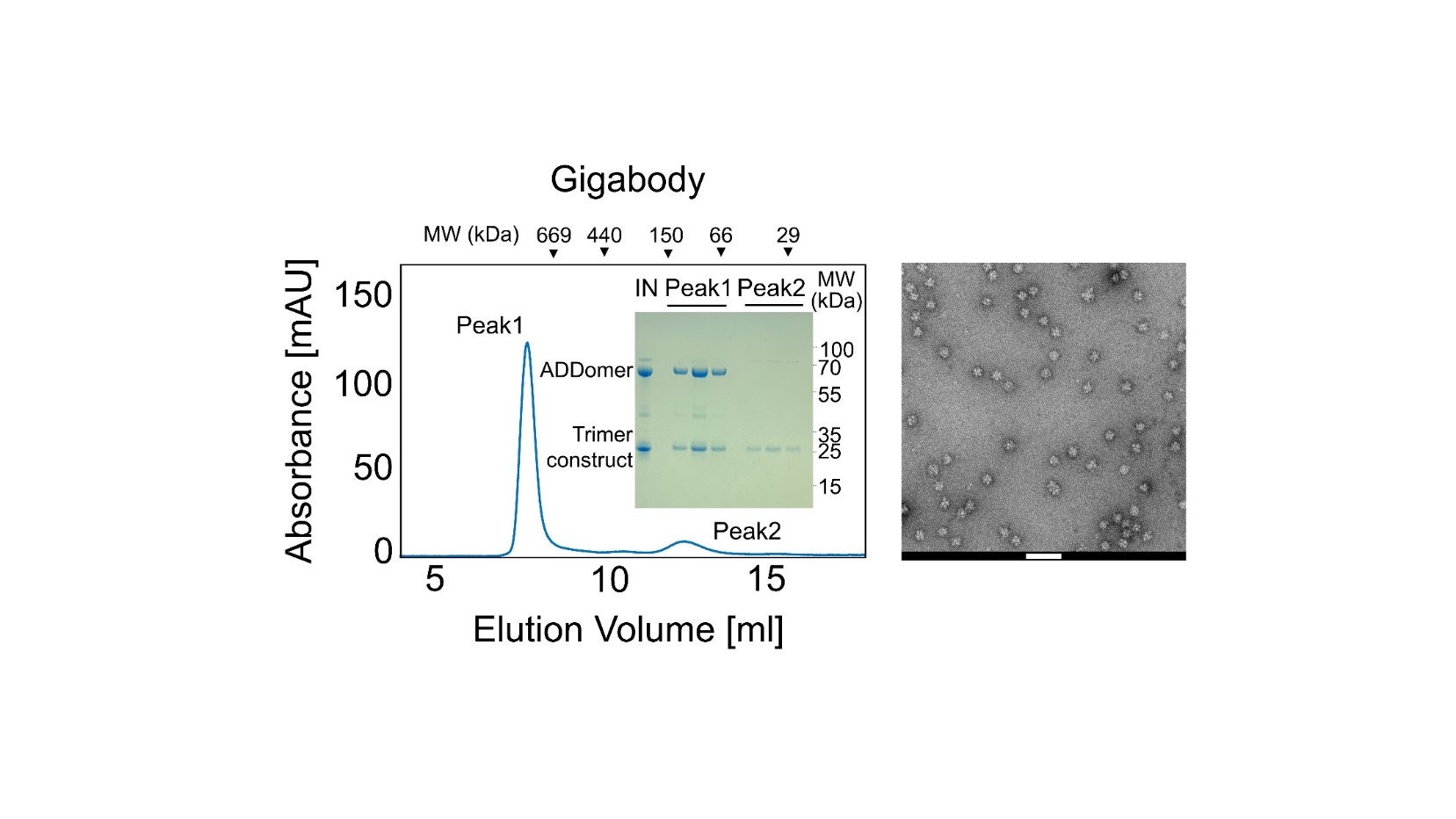

**Supplementary Fig. 8: Gigabody purification and quality control.** SEC of reconstituted Gigabody is shown (left). Molecular weight standard elution volumes are indicated (top). Gigabody elutes in peak 1 close to the void volume. Coomassie-stained SDS gel showing input fraction, peak 1 fractions with nanobody-trimer construct and penton base proteins and peak 2 fractions containing nanobody only are shown in the inset. Negative stain EM of peak 1 is shown on the right. Scale bar corresponds to 100 nm.

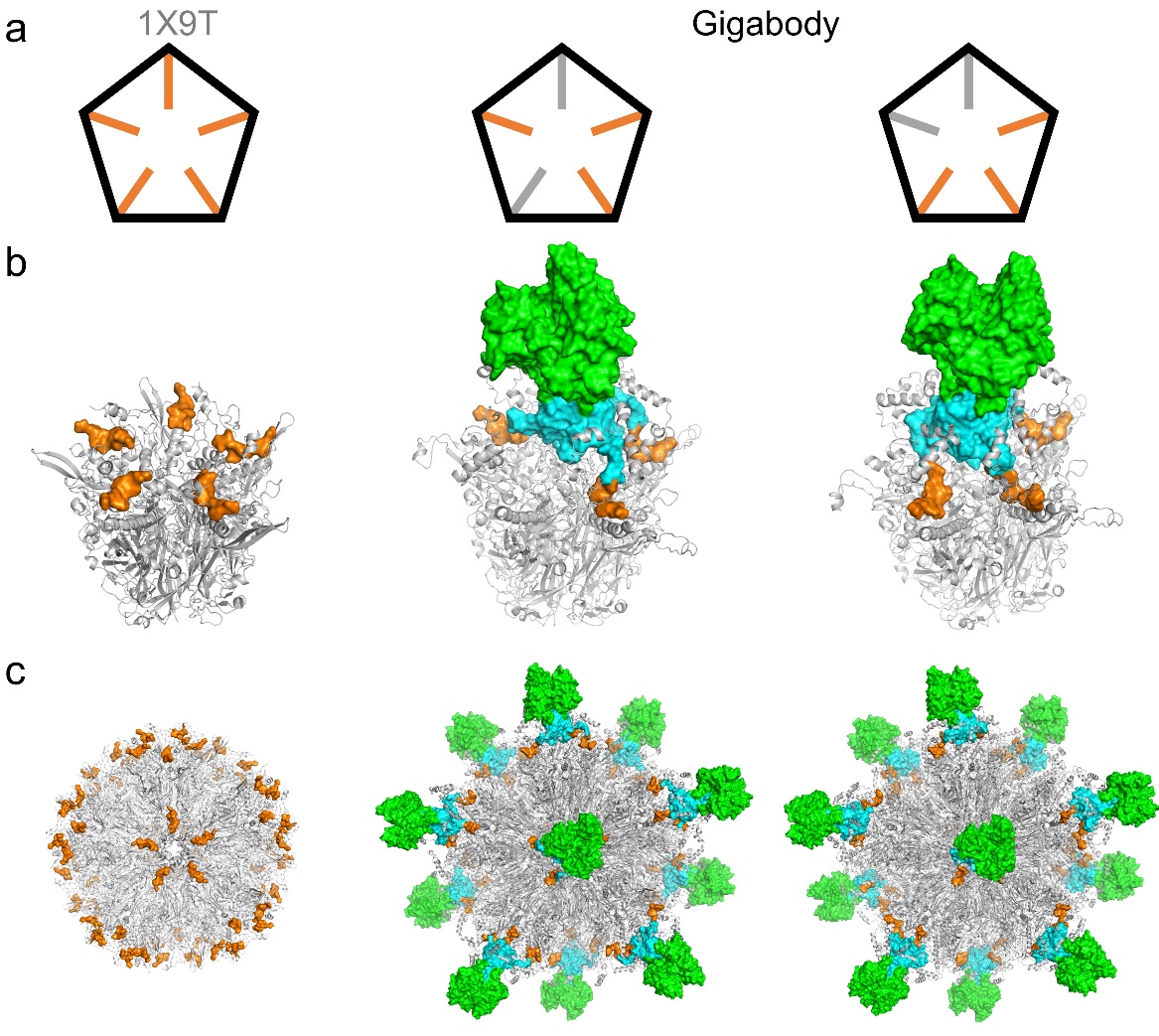

**Supplementary Fig. 9: Gigabody modeling.** Two distinct Gigabody models were constructed based on the Ad5 penton base fiber tail peptide complex structure (left, PDBID: 1X9T) ^1^ with the fiber tails (orange) either maximally spaced (middle) or attached to adjacent penton base protomers on the penton (right). **(a)** Illustration of the observed binding modes of the fiber tail peptides (orange) on the pentons (circumference drawn in black). Instead of natural three fiber tail peptides of the trimeric fiber protein, five peptides are observed in the crystal structure occupying all possible binding sites. In Gigabody, due to ADAH11 trimer binding, two of the binding sites remain unoccupied (grey). **(b)** Model of the complex of the chimpanzee adenovirus derived ADDomer penton (grey) and the trimer formed by ADAH11 nanobody (green) linked to T4 foldon trimerization domain (blue) and fiber tail peptide (orange). **(c)** Model of the assembled 60mer, constructed by aligning the pentons to their position in the experimental Ad5 fiber tail complex structure.

**Supplementary Table 1:**

Cryo-EM data collection and refinement statistics, ADDoCoV

| **Data Collection and Processing** | |
| --- | --- |
|  | ADDoCoV |
| Voltage (Kv) | 200 |
| Pixel size (Å/pix) | 1.05 (0.525) |
| Nominal magnification | 130,000 |
| Exposure (e- /Å2) | 44.03 |
| Frames per exposure | 40 |
| Defocus range (µM) | -0.7 to -2.2 |
| Final Particle Images | 32,227 |
| Symmetry imposed | I4 |
| Resolution at FSC = 0.143 (Å) | 2.36 |
| Map-sharpening B factor (Å^2^) | -25 |
| **Atomic Model** | |
| **Model composition** |  |
| Non-hydrogen atoms | 211740 |
| Protein residues | 25980 |
| **RMS deviations** |  |
| Bond lengths (Å) | 0.011 |
| Angles | 1.298 |
| **Validation** |  |
| MolProbity score | 1.53 |
| Clashscore | 1.93 |
| Rotamer outliers (%) | 0 |
| C-beta deviations | 0 |
| **Ramachandran plot** |  |
| Favoured (%) | 89.04 |
| Allowed (%) | 10.73 |
| Outlier (%) | 0.23 |

**Supplementary Table 2:**

Sequences of nanobodies selected by Ribosome display.

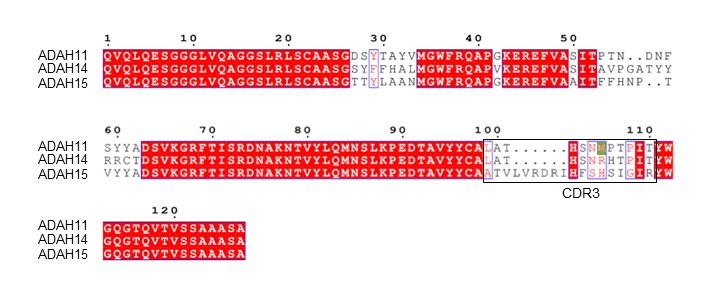

*CDR3 boxed in black, ADAH11 R105 underlaid in green.*

**Supplementary Table 3:**

Cryo-EM data collection statistics, ADDoCoV-ADAH11 complex

| **Data Collection and processing** | |
| --- | --- |
|  | ADDoCoV-ADAH11 |
| Voltage (Kv) | 200 |
| Pixel size (Å/pix) | 1.05 (0.525) |
| Nominal magnification | 130,000 |
| Exposure (e- /Å2) | 55.6 |
| Frames per exposure | 45 |
| Defocus range (µM) | -0.8 to -2.0 |
| Final Particle Images | 13950 |
| Symmetry imposed | C1 |
| Resolution at FSC = 0.143 (Å) | 4.06 |
| Map-sharpening B factor (Å^2^) | -79.66 |

**Supplementary Table 4:**

Adenoviral fiber tail peptide sequences

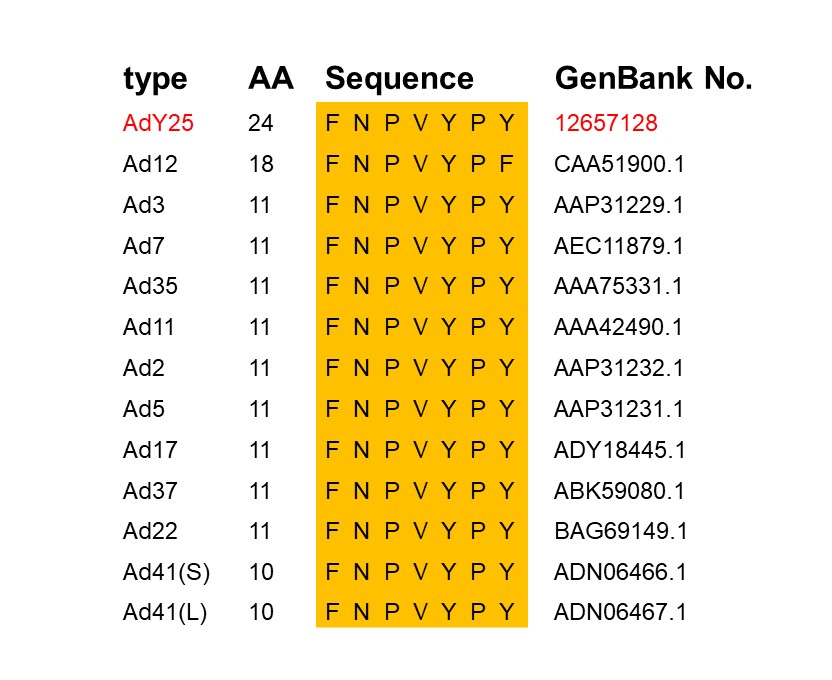

**Supplementary Table 5**:

Sequences of penton base protomers used in this study

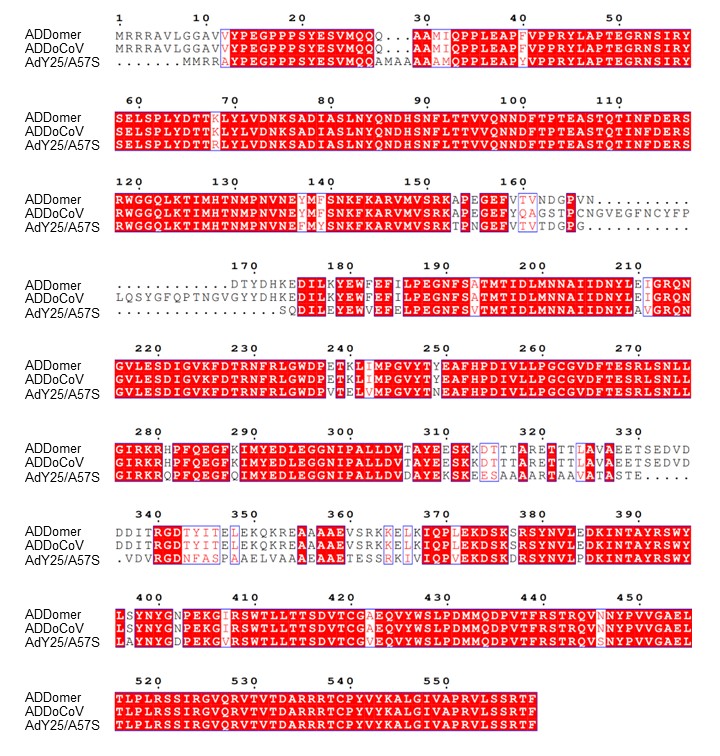

**Supplementary Table 6:** Sequences of proteins used for Gigabody preparation.

| **ID** | **Primary sequence** |
| --- | --- |
| AdY25 ADDomer A57S protomer | MMRRAYPEGPPPSYESVMQQAMAAAAAMQPPLEAPYVPPRYLAPTEGRNSIRYSELSPLYDTTRLYLVDNKSADIASLNYQNDHSNFLTTVVQNNDFTPTEASTQTINFDERSRWGGQLKTIMHTNMPNVNEFMYSNKFKARVMVSRKTPNGEFVTVTDGPGSQDILEYEWVEFELPEGNFSVTMTIDLMNNAIIDNYLAVGRQNGVLESDIGVKFDTRNFRLGWDPVTELVMPGVYTNEAFHPDIVLLPGCGVDFTESRLSNLLGIRKRQPFQEGFQIMYEDLEGGNIPALLDVDAYEKSKEESAAAARTAAVATASTEVDVRGDNFASPAAELVAAAEAAETESSRKIVIQPVEKDSKDRSYNVLPDKINTAYRSWYLAYNYGDPEKGVRSWTLLTTSDVTCGVEQVYWSLPDMMQDPVTFRSTRQVSNYPVVGAELLPVYSKSFFNEQAVYSQQLRAFTSLTHVFNRFPENQILVRPPAPTITTVSENVPALTDHGTLPLRSSIRGVQRVTVTDARRRTCPYVYKALGIVAPRVLSSRTF |
| ADAH11-Trimer | MKYLLPTAAAGLLLLAAQPAMAQVSKKRARVDDTFNPVYPYDADNAPTVPFINPPFVSSDGFQEKPSGRLVPRGSPGSGYIPEAPRDGQAYVRKDGEWVLLSTFLGGGSQVQLQESGGGLVQAGGSLRLSCAASGDSYTAYVMGWFRQAPGKEREFVASITPTNDNFSYYADSVKGRFTISRDNAKNTVYLQMNSLKPEDTAVYYCALATHSNRPTPITYWGQGTQVTVSSAAASAHHHHHHKLDYKDHDGDYKDHDIDYKDDDDK |
| *Fiber peptide underlaid in orange, T4 foldon in blue, ADAH11 nanobody in green.* | |

**Supplementary Movie 1: Architecture of ADDoCoV nanoparticle vaccine**. This movie shows ADDoCoV based on cryo-EM data and MD simulations. SARS-CoV-2 RBM derived epitopes (60 copies per ADDoCoV) are coloured in red. Pentons forming the nanoparticle scaffold are coloured in shades of cyan, lilac and gray.

**Supplementary Movie 2: Gigabody nanoparticle displaying ADAH11 nanobody trimers.** This movie shows the Gigabody nanoparticle comprising 12 trimers of ADAh11 nanobodies (coloured in green) fused to a T4 foldon trimerization domain (blue) and an Adenovirus AD25Y fiber tail peptide (orange). Pentons are colored in shades of cyan, lilac and gray.
